# A spectral jamming avoidance response does not help bats deal with jamming

**DOI:** 10.1101/2019.12.16.876086

**Authors:** Omer Mazar, Yossi Yovel

## Abstract

For decades, researchers have speculated how echolocating bats deal with acoustic interference created by conspecifics when flying in aggregations. It is thus surprising that there has been no attempt to quantify what are the chances of being jammed, or how such jamming would affect a bat’s hunting. To test this, we developed a computer model, simulating numerous bats foraging in proximity. We used a comprehensive sensorimotor model of a hunting bat, taking into consideration the physics of sound propagation and bats’ hearing physiology. We analyzed the instantaneous acoustic signals received by each bat, and were able to tease apart the effects of acoustic interference and of direct resource competition. Specifically, we examined the effectiveness of the spectral Jamming Avoidance Response - a shift in signal frequencies - which has been suggested as a solution for the jamming problem. As expected, we found that hunting performance deteriorates when bats forage near conspecific. However, applying a Jamming Avoidance Response did not improve hunting, and our simulations clearly demonstrate the reason: bats have adequate natural signal variability due to their constant adjustment of echolocation signals to the task. The probability to be jammed is thus small and further shifting the frequencies does not mitigate spectral jamming. Our simulations reveal both negative and positive insight: they show how bats can hunt successfully in a group despite potential sensory interference and they suggest that a Jamming Avoidance Response is not useful.

## Introduction

Echolocation, a prime example of active sensing, provides bats with the ability to detect and hunt flying insects while avoiding obstacles in total darkness^1^. Echolocating bats emit high-frequency sound-signals and process the reflected echoes to sense their surroundings. While hunting in a group, conspecific bats emitting signals with similar frequencies may interfere with the ability of nearby bats to detect and process their own echoes. Understanding how bats avoid this interference, which is referred to as ‘jamming’ or ‘masking’, and how they segregate the desired weak echoes from the much louder calls emitted by other bats is one of the most central debates in the field. We define a ‘masking signal’ as any signal that may interfere with the bat’s ability to detect and localize an echo, and a ‘jamming signal’ as a signal that completely blocks the detection of an echo (see Methods).

The question of how bats deal with conspecific masking and whether they perform a spectral Jamming Avoidance Response (JAR) has been widely studied but is still under dispute. Many studies have suggested that bats change their echolocation frequencies when hunting the presence of other bats^2–5^ or when exposed to playback partially or fully overlapping signals^6–9 2–8, 10, 11^. Particularly, in this study, we only deal with *spectral* JAR (which we will term JAR). In contrast, several recent field-studies and laboratory experiments found no evidence for a use of spectral JAR by bats^12–14^.

The main goal of our study is to use a mathematical approach in order to deepen the understanding of the masking problem and its impact on bats’ hunting, and specifically to examine whether an intentional shift of signal frequencies (i.e., a spectral JAR) can assist bats to mitigate the masking problem. We developed an integrated sensorimotor model of bats pursuing prey. The modeling approach entails several advantages in comparison to studies with real bats. (1) It allows us to assess the exact acoustic input received by each of the hunting bats at every instance. This is currently impossible to do in reality even when a microphone is placed on the bat. (2) Modeling enables manipulation of different parameters and examining their influence on masking including testing hypothetical scenarios that tease apart factors that are coupled in reality.

We analyzed the effect of masking under various prey and bat densities and when using different echolocation behaviors. We measured the probability of jamming, the hunting performance and we explicitly examined whether applying a spectral JAR improves hunting performance when hunting with conspecifics. We were able to discriminate between the effect of direct interference resulting from the need to avoid conspecifics and to compete with them over prey, and the effect of sensory masking due to conspecific calling. We show that shifting the emission frequencies (i.e., a JAR) does not assist mitigating masking because bats’ signals already differ from each other due to their well-known behavior of adjusting their signals based on the task and the environment.

## Results

The model consists of numerous bats searching for and attacking prey in a confined 2D area using echolocation. Each simulated bat transmits sound signals and receives the echoes returning from prey items, as well as the signals emitted by conspecifics, which might mask or jam its own echoes. The prey’s movement mimics a moth^21, 22^ with no ability to hear the bats. Prey echoes are detected and localized based on biological-relevant assumptions which consider sound reflection and propagation, and hearing physiology (see Methods). Based on the incoming acoustic information, the bat decides whether to continue searching, to pursue prey or to avoid obstacles such as other bats. It then adjusts its echolocation and movement according to the vast literature on bat echolocation ^1, 23^, and the recently published control models of bat flight and hunting^15, 16, 24–28^. For example, the simulated bats emit search signals with a power of 110 dB-SPL (at 0.1m) and they lower their power (and adjust other echolocation parameters) when approaching prey. A successful hunt (i.e. a capture) occurs only when the simulated bat gets within 5cm from the prey. That is, the bats sometimes initiate attacks but miss.

We first demonstrate that our simulations behave similarly to bats. The simulated bats managed to detect, pursue and capture prey at high rates both when hunting alone and when hunting in a group (see Figure 1 and Movie S1 for examples of hunting by simulated bats). The movement parameters of the bats in both single and multiple individual scenarios were similar to those of actual bats, suggesting that our model managed to capture the essence of the foraging movement (Fig S 1).

**Figure 1:**
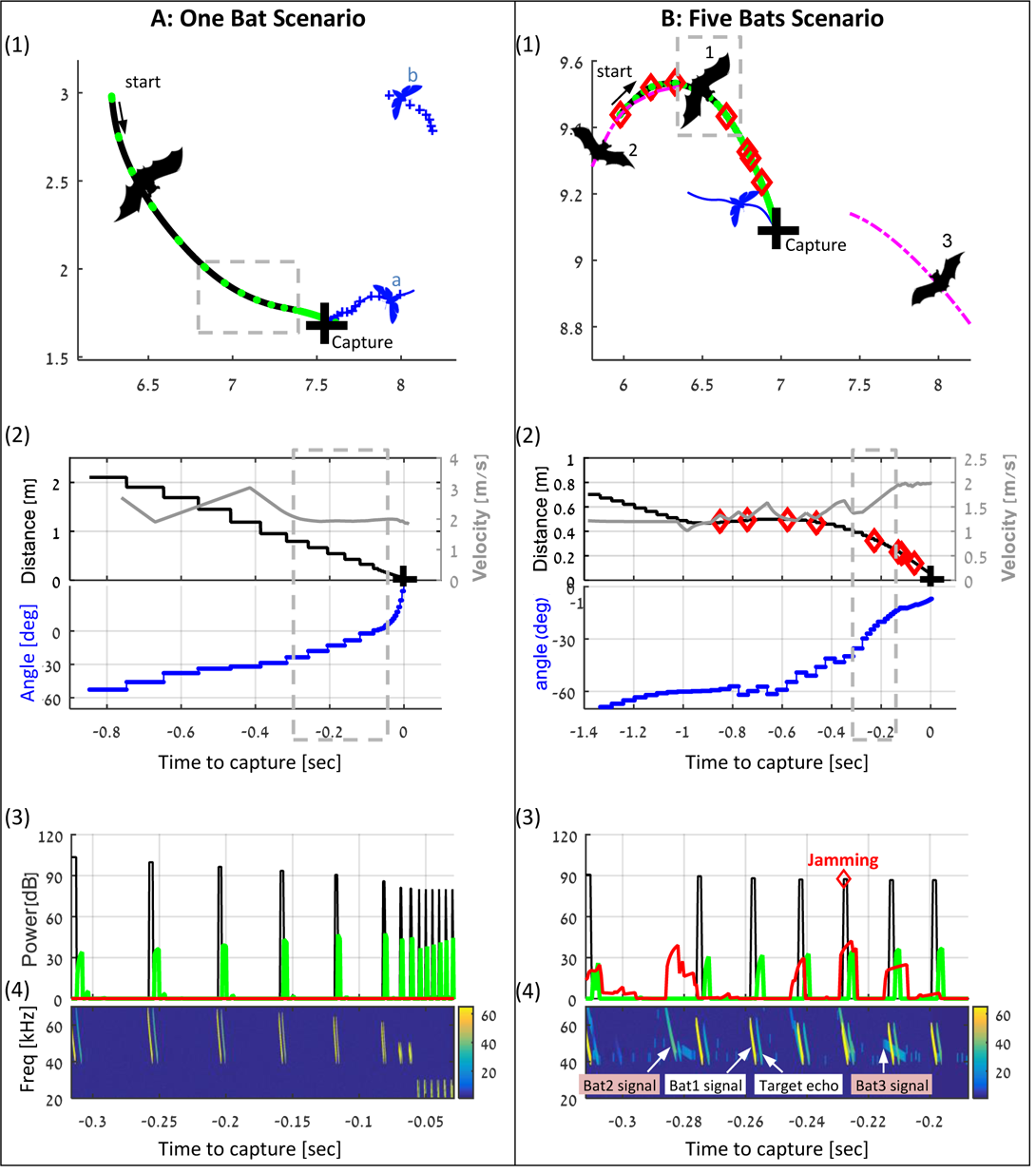
Examples of individual and group hunting of the simulated bats (also see supplementary movie). (A) A single simulated bat flying in the arena with 5 prey items. The bat detects two prey items and decides to pursue the closest one (moth a). **Panel 1** shows the trajectories of the bat (black) and the moths (blue), the positions of each emitted signal (green dots), and the location of the capture (black cross). **Panel 2** shows the bat’s velocity (gray), the distance (black) and relative angle (blue) to target. **Panel 3** shows the bat’s received acoustic scene for the section marked by a gray dashed line, including the envelopes of the following signals: the transmitted signals (black), the received prey echoes (green), and the received masking signals (red). **Panel 4** shows a spectrogram of the same segment as in panel 3. Note how the signals’ frequency drops at the final terminal buzz. **(B)** Three bats (out of five in the arena) hunting in an environment with 10 moths. Bat 1 detects and pursues a moth, while conspecifics (bats 2 and 3) fly nearby, emitting echolocation signals some of which mask (or jam) the echoes received by bat 1. Instances of jamming are marked with red diamonds in panels B1-4. All colors and symbols in B1-4 are the same as in A1-4. Magenta lines depict trajectories of conspecifics. In panel B4 we depict one signal for each bat, illustrating the variations between the signals due to the different behavioral phases of each bat. Note that bat 2 also detects the same prey item and pursues it and thus its signals are ‘approach’ signals. From detection to capture, bat 1 emitted 64 echolocation signals, 8 of which were jammed.

### The influence of a spectral JAR

we next compared the detection and localization performance of bats applying a JAR to bats that do not actively react to masking signals. In both groups, the frequencies of the individuals were sampled from a normal distribution with a standard deviation of 4 kHz, as observed in nature^15, 16, 29^, and they adjusted their signals according to their task and their distance to object. In some simulations, the bats applied a JAR, namely, whenever their signals were jammed – they shifted their terminal frequency away from the jamming signal in steps of 2kHz and they kept shifting the frequency as long as masking occurs (the signal was shifted either upward or downward in the opposite direction of the jamming frequency, Methods). The entire frequency range of the signals was shifted upward or downward. We tested two different receiver models with different assumptions: 1) the ‘correlation-detector’ which is an optimal receiver and is at least slightly better than the bat’s brain^30–35^; (2) the ‘filter-bank receiver’ which is considered to represent the mammalian auditory physiology and implements a series of gammatone band-pass filters^34,36–39^ (see Methods).

The main function of a spectral JAR should be to improve detection by decreasing the overlap between the spectra of the masking signal and the echo. We, therefore, start with examining whether a JAR under different conditions (e.g., different receiver models and different bat densities) achieves this goal. To quantify this, we used four detection-criteria (see Methods): (1) The jamming-probability is defined as the probability that the echo reflected from the closest prey item is totally jammed by a masking signal and is thus not detected. (2) The SNR (Signal to Noise and interference Ratio) is defined as the ratio between the peak intensity of the detected echo, and the peak intensity of all masking signals. (3) The ranging error is the difference between the estimated and the actual distance to prey. (4) The false-alarm rate is the probability of identifying a masking signal as prey by mistake.

In all scenarios, and for the two different receiver models, the jamming avoidance response did not decrease spectral masking (see Fig S 2), and did not improve detection performance according to any of the four criteria defined above. See Figure 2: One-way ANOVA statistics for correlation and filter-bank, respectively**: jamming-probability**: F_1,237_= 2.96, p=0.09; F_1,237_=2.26, p=0.13, **SNR:** F_1,237_=0.62, p = 0.43; F_1,237_= 0.02 p=0.88, **ranging error:** F_1,237_= 0.1, p =0.76; F_1,241_= 0.01, p=0.96, **false-alarm:** F_1,237_=0.19, p=0.66, for filter-bank only. A possible explanation for this seemingly surprising result is the fact that a bat’s signals continuously vary depending on its behavioral phase and its distance to the targets. Therefore, at any instance, the signals of two bats will already differ, even if their signal repertoire is identical. Therefore, the influence of additional variability between the signals, achieved by spectral JAR, is insignificant, see Fig S 2.

**Figure 2:**
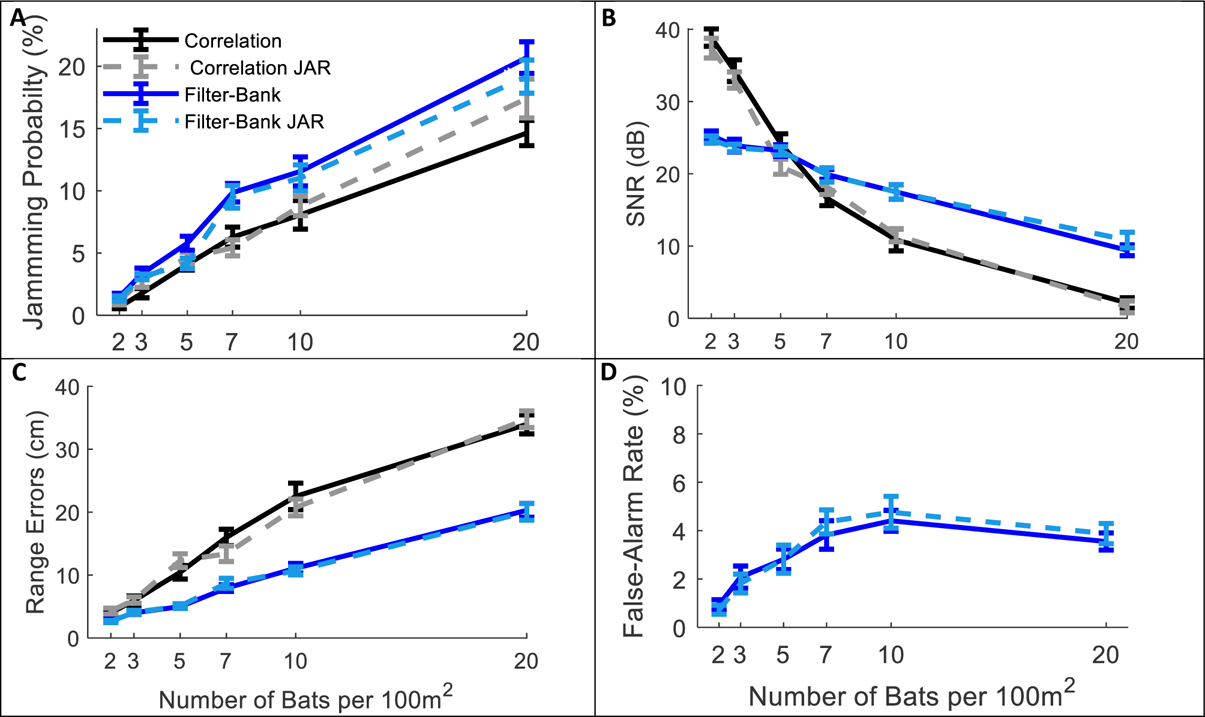
Applying a JAR does not improve detection. Panels A-D depict four detection criteria as a function of bat density for a correlation-receiver (solid black), and a filter-bank receiver (solid blue) with and without a JAR (dashed gray and dashed turquoise respectively). The prey density was constant with 20 prey items per 100m^2^. Error-bars depict standard-errors in all panels. Each data point represents 60-80 simulated bats. **(A)** Jamming Probability. **(B)** SNR-The SNR is the Signal to Interference Ratio, measured on all detected prey items. **(C)** Range errors. Note that the range-errors are high because they are calculated for all detected prey. The errors for the pursued prey (i.e. prey items that bats attacked) are substantially lower, since, as the bats approach their targets echoes have higher SNR and the errors decrease. **(D)** False-alarm rate, which is measured only for the filter-bank model.

### The effect of masking on hunting

next, we tested the effect of masking on hunting performance (i.e., the prey capture rate) under different scenarios. We started with a hypothetical scenario (‘no-masking’), in which bats forage in a group without any masking, i.e. they detect and pursue prey regardless of any sensory interference, but they still have to avoid other bats and sometimes lose prey due to competition. This null-hypothesis scenario enabled us to estimate the non-sensory effects of group hunting, which would be very difficult to do in an experiment with real bats. Ultimately, it allowed us to isolate the effect of sensory interference only. Hunting performance was measured in different bat-densities (from 1 to 20 bats per 100m^2^) and different prey densities (3, 10, 20 moths per 100m^2^), see Figure 3.

**Figure 3:**
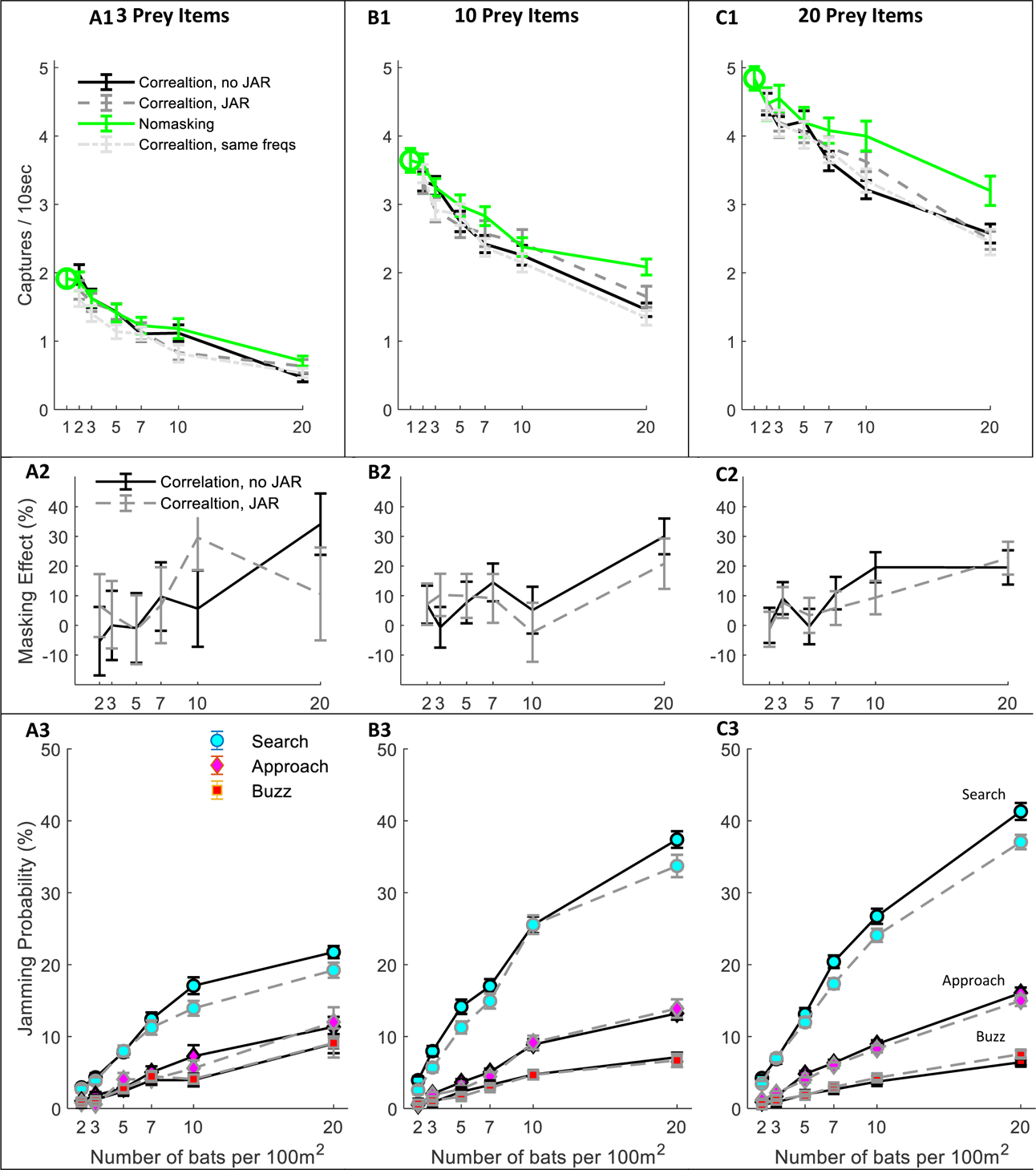
Hunting Performance for different receiving models. The hunting success rate is presented for three prey densities per 100m^2^: **(A)** 3 prey items**; (B)** 10 prey items; **(C)** 20 prey items. Line colors and styles depict the performance of different receiver models: green - no-masking; solid black - correlation-detector with random signal variability; dashed dark gray - correlation-detector with JAR response; dash-sotted light gray - correlation-detector without signal variability. Panels **(A1-C1)** depict the performance as a function of bat density; a green circle shows the performance of a single bat. The regression slopes of no masking condition (green lines) are (mean± SD): 0.06± 0.0065 0.08± 0.0084, 0.079± 0.013 captures per bat per ten seconds, at the prey densities above, respectively (ANCOVA: F_1, 564_ = 84.3, p<0.0001; F_1, 679_ = 90.4, p<0.0001; F_1, 431_ = 37.8, p<0.0001). There was no significant difference in performance when applying or not applying spectral JAR - see main text and compare gray and black lines. Panels **(A2-C2)** show the masking-effect on hunting, i.e., the relative decrease in hunting relative to the ‘no-masking’ condition. **(A3-C3)** The probability of jamming at different behavioral phases: search (turquoise marker), approach (magenta marker) and buzz (red markers). Jamming probabilities during the search were significantly lower by at most 4.5% when using a JAR (ANCOVA, F_1, 2422_=23.42, p<0.0001). However, in the approach and buzz phases (which are more critical for foraging), there was no significant difference between the two models (ANCOVA, F_1,2388_=0.11, p=0.74; F_1, 2347_ = 0.11, p=0.73, respectively). Error-bars show means and standard-errors for 70-120 simulated bats in each data-point.

Even without any sensory interference (i.e., in the ‘no-masking’ condition), hunting performance significantly degraded as bat density increased due to competition over prey and due to the need to avoid conspecifics (Figure 3A-C; see Green lines). The reduction in performance was significant in all prey densities and resulted in a maximum decrease of 36%, 57% and 67% in performance when the bats’ density increased (from 1 to 20) at three prey densities: 3, 10 and 20 prey items per 100m^2^, respectively. See Figure 3 A1-C1, One-way ANOVA, F_1, 188_ = 64.3, p<0.0001; F_1, 188_ = 58.9, p<0.0001; F_1, 128_ = 36.1, p<0.0001, respectively.

We next examined the masking-effect which we defined as the reduction in performance resulting from sensory masking only, relative to the no-masking performance (Equation 1).

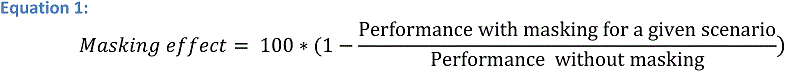

Sensory masking further hindered hunting under all conditions, but there was no significant difference in performance whether the bats used a JAR or not (Figure 3, A1-C1, compare blue, black and magenta lines; ANCOVA, F_3, 1218_ = 2.53, P=0.08, F_1, 1581_ =0.57, P=0.56, and F_1, 1085_ =0.24, P= 0.78, for 3, 10 and 20 prey items per 100 m^2^, respectively. This was the case also for another theoretical scenario that we tested: the ‘no-frequency variation scenario’, in which all bats had the same signal repertoire and, hence, it simulates an extreme case with maximal sensory interference (One-way ANOVA, between ‘no-frequency variation’ and JAR with random frequencies: F_2,856_=2.74, p=0.1; F_2,971_=0.85, p=0.36; F_2,1086_=0.43, p=0.64, for prey densities of 3,10 and 20 prey items per 100m^2^, respectively). See Fig S 3 for a similar analysis for the filter-bank model.

To deepen our understanding of why a JAR does not improve performance, we analyzed the jamming-probability in different behavioral phases. We found that jamming mostly occurred during the search phase while, as the bats shifted from the search to the approach phase, the probability of jamming decreased significantly because the prey’s echoes become louder (Figure 3 A3-B3-C3, ANCOVA, for the comparison between search and approach at any bat and prey density: F_1, 8527_ =2784, p<0.0001, panels B3-C3 show that less than 15% of the prey echoes are jammed during the approach phase). Notably, if a potential prey echo is jammed in the search-phase, the bat is likely to detect the prey with one of its following emissions, so jamming during the search phase is less detrimental for foraging. The low probability of jamming during the approach is probably the main reason for the relatively small effect of sensory masking on performance.

We also analyzed the causes of unsuccessful attacks when bats initiated an attack but failed to capture prey. There were four reasons for failed attacks: avoiding a nearby conspecific, losing the prey to a conspecific, avoiding an obstacle (the borders of the arena) and missing the prey due to an insufficient maneuver or due to sensory error (resulting for example from jamming). We analyzed the proportion of these different sources of failure with and without sensory masking (see Fig S4). With 20 bats and 10 prey items per 100m^2^, without masking, 34%±2% of the capture attempts were successful (mean±SE). The unsuccessful attempts were due to conspecifics avoidance: 27%±2%; lost prey to conspecifics: 17%±1.5%; obstacle avoidance: 7%±2% and misses: 15%±2%. When sensory masking was added, the proportion of successful captures significantly decreased to 26%±2% (One-way ANOVA, F_1, 198_=4.59, p=0.033), and misses became the most substantial cause for failure, significantly increasing to 38%±3.5% (One-way ANOVA: F_1, 198_=68.8, p<0.0001). The total number of hunting attempts, however, was not affected by the masking (effect-size=0.01 trials per 10 seconds; one-way ANOVA: F_1, 198_=0.08, p=0.97).

### The effect of prey density on group hunting

Finding and catching prey is easier when prey is abundant, and as excepted, the hunting rate significantly increased as a function of prey density in all bat densities see Figure 4A. The regression slopes for the correlation-detector indicates an improvement of 0.09±0.001, 0.011±0.001, 0.086±0.001, 0.08±0.001 captures per additional prey item per 10 seconds, for 1, 5, 10 and 20 bats per 100 m^2^, respectively; mean± SD are reported. The masking effect did not change significantly as a function of prey density, see Figure 4B, Pearson Regression: F_2, 48_ = 1.2, p=0.28. Assuming that a bat arriving at a foraging site can roughly estimate the density of prey and the density of bats in the area, we examined how the ratio of these two densities affects performance. Hunting performance drops rapidly with an increase in the bat to prey density ratio, but that it levels out once the bat to prey ratio increases above ‘1’. In general, the same pattern was observed for different bat densities (Figure 4C), but for a given bat to prey ratio, the performance was better when there were more bats (and more prey). The improvement in performance as a function of bat density, when the bat-to-prey ratio was below 1, was significant in scenarios with masking (ANCOVA: F_1, 1093_ = 38.2, p<0.001).

**Figure 4:**
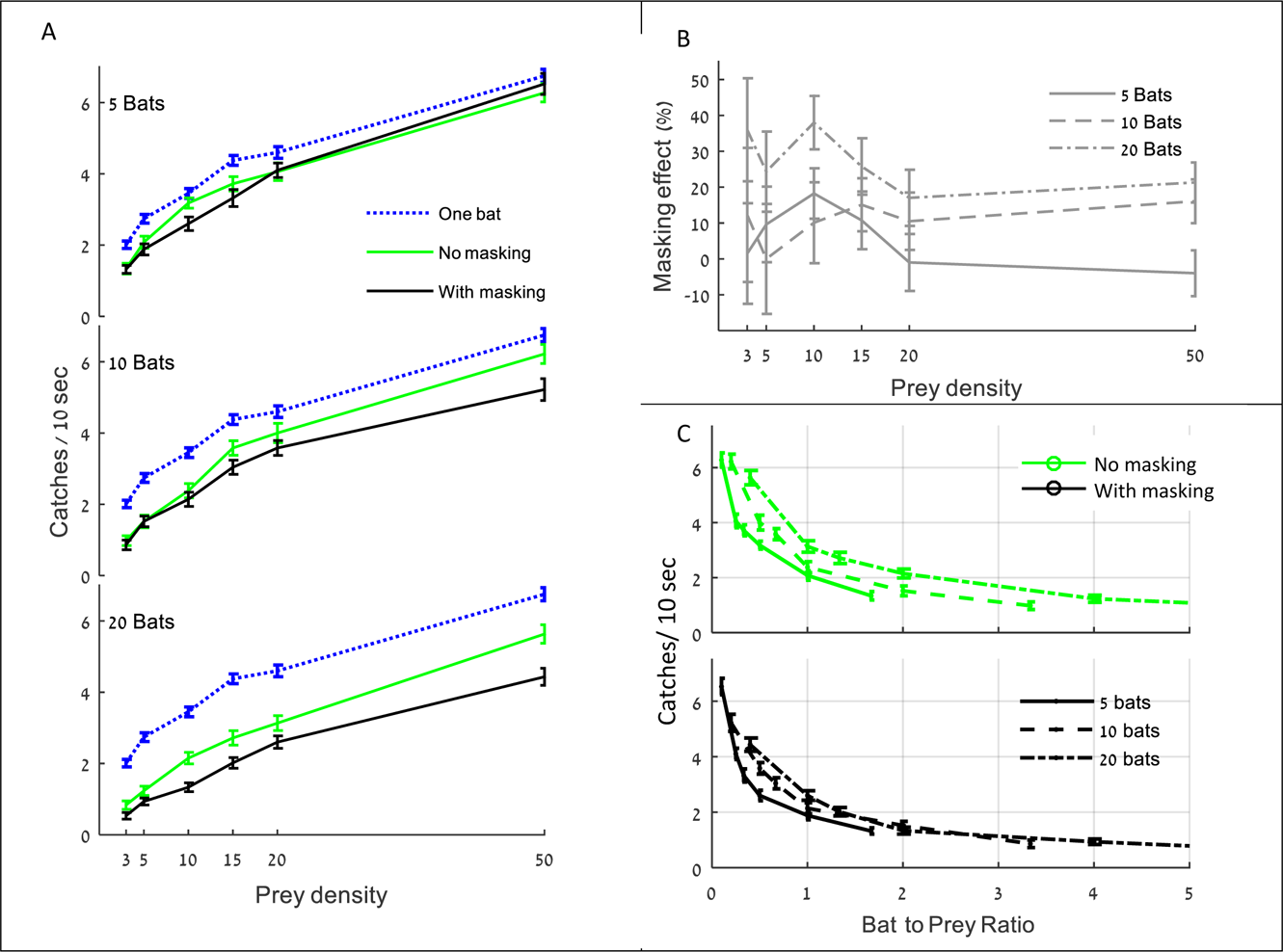
The influence of prey density on performance. (A) Hunting rate vs. prey density for three different bat-densities for the following scenarios: ‘no-masking’ (dashed line), masking (solid line), and a single bat as a reference (dotted blue line). Error bars represent means and standard-errors for 50-100 simulations in each data-point. **(B**) The masking effect for the same scenarios as in (A). The masking effect was computed according to Equation 1. **(C)** Hunting performance as a function of the proportion between bat density and prey density. The upper panel depicts performance in the ‘no masking’ (green) condition and the lower panel shows the performance with masking (black), for three bat densities. In all panels, results ‘with masking’ are shown for the correlation-detector with signal frequencies distributed normally with a mean of 39 kHz and SD of 4 kHz.

### The effect of the echolocation signal design and the detection threshold on hunting in a group

after observing that spectral changes do not assist mitigating jamming, we tested whether other adjustments to echolocation or physiological parameters could improve bats’ performance when hunting in a group. We tested the effect of two prominent signal parameters: power and duration, and we also examined the influence of the detection threshold, which is a function of the auditory system (for all parameters, we used a range of values suggested in the bat literature). For each of these parameters, we first examined its effect on the overall hunting success when foraging in a group (i.e., including both direct competition and sensory interference), and we then examined the parameter’s effect specifically on the masking.

Increasing the signal’s power improved hunting performance, but only up to power of ca. 110-120 dB-SPL, (at 0.1m) above which the improvement was negligible and insignificant (Figure 5 A1, shows that performance increased significantly when signal power increased from 90 to 110 dB: ANCOVA, slope=0.06 captures/dB, F_4, 3815_=386, p<0.001; performance did not change when signal power increased from 120 to 150dB: F_4, 4795_=0.1, p=0.98). Interestingly, the power of the signal had no effect on the masking effect (i.e., the reduction in performance beyond the no-masking condition), suggesting that increasing the emission power assists hunting in general (through increasing the detection range) and does not assist overcoming the masking problem specifically (Figure 5A2; F=2.45, p=0.16, Pearson linear regression).

**Figure 5:**
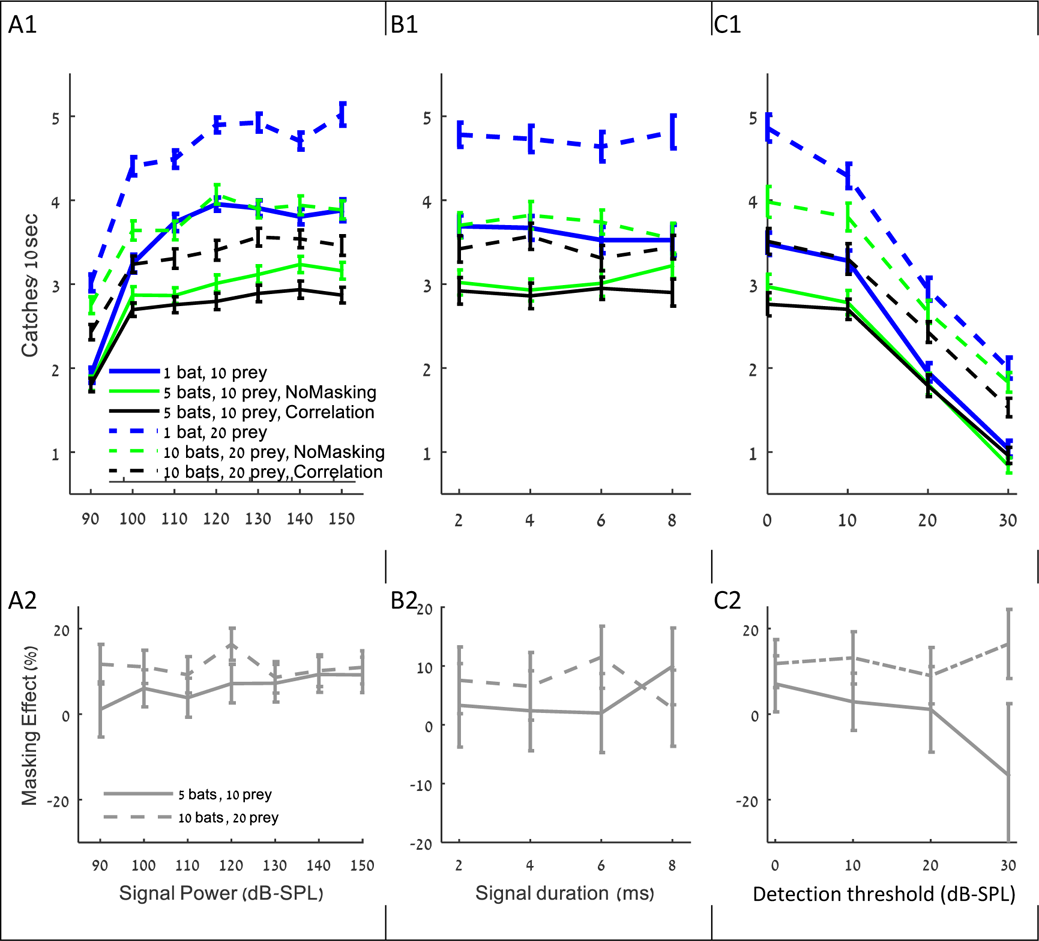
The influence of (A) signal power, (B) detection threshold and (C) signal duration on the performance (A1-C1), and masking effect (A2-C2). The panels include the following conditions: the correlation-detector (with bat frequencies distributed normally, case 2 above, black), no-masking (green), and one bat (blue). Solid lines indicate a density of 10 prey items while dashed lines represent scenarios with 20 prey.

Changing the duration of search-signals had no significant effect on the hunting performance (Figure 5 B1: ANCOVA, F_1, 2314_=0.15, p=0.69). To test the effect of signal duration we slightly varied the simulation (see the detection section in the Methods).

Decreasing the hearing threshold (i.e., increasing hearing sensitivity) significantly improved hunting (Figure 5 C1, ANCOVA, F_4, 2395_=915, p<0.0001). Like in the case of increasing emission power, changing the hearing threshold did not significantly change the masking effect (Figure 5 C2; ANCOVA, F_1, 797_=2.19, p=0.14). This is probably because the decrease in the hearing threshold increases the probability of detecting both echoes and masking signals. With a high hearing threshold of 30dB and a density of 5 bats per 100m^2^, there seems to be a negative masking effect, but that is because prey is only detected from very short distances and thus prey detection and masking hardly occur, and consequently, the standard error of our estimate under these conditions is high.

## Discussion

The jamming problem is one of the most fundamental problems raised by researchers of echolocation, but to our best knowledge, nobody ever estimated what are the chances of being jammed by another bat and how such jamming would actually affect hunting performance. This is very difficult to do with real bats as even if a microphone is placed on the bat, it is typically not as sensitive as the bat itself and it is not placed inside the ear. The substantial body of literature that has accumulated on bat echolocation and sensorimotor control now allows simulating natural scenarios where bats are foraging in aggregations. Using this approach, we show that even in very high bat-densities, bats can probably capture insects at high rates. Because we always underestimated the bats’ performance (see Methods), this result probably reflects reality too. Notably, we did not fit any of the model’s parameters – we used parameters that are based on our measurements on real bats or published results. Similarly, we used a simple control strategy to steer the bat to the prey. For example, we do not assume any memory of the position of the target while real bats probably use memory to overcome temporal miss-detections caused by momentary jamming. The ability of real bats to catch prey and avoid obstacles under severe masking was corroborated by several behavioral studies^10, 13, 23, 31, 40^.

Furthermore, our analysis is based on relative measurements between different scenarios, therefore, even if the exact rates of prey-capture that we estimated are biased somehow, the principles which we observed are likely correct, providing insight to the jamming problem. The two most important results are: (1) Much of the interference that bats suffer from when foraging in a group results from competition over prey and from the need to avoid conspecifics, and not from acoustic masking. One of the main reasons for this is that jamming mostly occurs when the bat is still searching for prey, while once it has detected prey and is closing in on it, prey echoes become loud and the chances of jamming substantially decrease. Masking during search might sound problematic, but even if a prey’s echo is completely jammed, another echo from the same prey will likely be detected with one of the following echolocation signals. (2) Using a spectral JAR, which has been suggested by many previous researchers, is ineffective for solving the jamming problem or even reducing it. The reason for this is that bats constantly change their signals according to behavioral phase and distance to nearby objects. Even if two bats have the same signal repertoire, at any moment in time, their signals are different due to the different behavioral phase they are in and because they are likely to have objects at different distances. Moreover, we only used the bats’ first harmonic. Simulating the second harmonic too, thus using signals with more than twice as much bandwidth, would have probably made the jamming avoidance response even more irrelevant, because the differences between the signals of different bats would be naturally larger at any moment (even without a JAR). In theory, in bats that emit narrow bandwidth signals, such as bats with shallow FM calls^41^, jamming might be more influential and a spectral JAR might be more beneficial. However, most shallow FM bats increase bandwidth considerably when pursuing prey, and thus spectral JAR is probably not substantial for those bats too, at least during the pursuit. Indeed, in a previous study, we did not observe a spectral JAR in a bat that uses shallow FM calls (*R. microphyllum*)^12^.

Both of the receiver models that we tested revealed the same results regarding the inefficiency of a JAR. One of the models (the correlation-receiver) is considered optimal in terms of its detection abilities and probably over-estimates bats’ abilities. The fact that such a detector, which is extremely sensitive to the specific spectro-temporal pattern of the desired signal, did not show better performance when a JAR was applied, strongly suggests that the JAR is not helpful for real bats too, as their ability to use the differences induced by a JAR are lesser, compared to this receiver.

As expected, the correlation-receiver outperforms the filter-bank in all scenarios; it has higher SNR, it detects more objects (see Fig S 5), and it has a lower probability to be jammed by masking signals (see Figure 2 a,b). Consequently, the total hunting performance is better (compare Figure 3 A1-A3 with Fig S 3 A1-A3). The range-errors of the filter-bank seem lower (Figure 2c), but this is only because the correlation-receiver detects farther objects than the filter-bank receiver and these objects have lower SNR and thus higher range errors. Apparently, this larger error has a little effect on the performance, as the bats get closer to targets, the SNR improves and the errors decrease.

Another interesting result of the simulations was revealed when testing which of the echolocation parameters would allow bats to perform best when hunting in aggregations. We found that the emission power actually used by real bats when hunting in a group (ca. 110-120 dB-SPL^24, 42, 43^) gave the best performance in the simulation. Increasing the emission power mainly helped increasing prey detection range and not overcoming masking - the masking effect was the same independently of the emission power. Moreover, increasing emission power beyond this level did not further improve hunting probably because when hunting in aggregations there is no benefit in detecting prey beyond a certain distance. Prey that is farther than this distance is very likely to be detected and caught by a closer bat. This result demonstrates how everyone can call louder and still benefit up to a certain degree, corroborating our and others’ previous observations ^2, 13, 44^. It would have been difficult to explain the benefit of everyone calling louder without a simulation (note that we did not consider the caloric cost of increasing the emission power which might further reduce actual calling power in reality).

Changing the signal’s duration did not affect the performance. A possible explanation is the fact that all simulated bats increased their signal duration. Therefore, the benefit of own longer signals is apparently balanced with the greater probability of overlapping with conspecific signals. Note that we are not saying that signal duration irrelevant for hunting in general, but only that it does not affect the ability to mitigate masking and does not improve hunting in a group.

Our simulations also provide insight on bats’ behavioral ecology. An analysis of the effect of the bat-to-prey density on hunting performance indicates that performance drops rapidly when the bat-to-prey density ratio increases above ‘1’, but that the effect then levels out. On the one hand, this implies that in most cases (when the ratio is below 1) a bat should join an aggregation of bats because such an aggregation usually implies the presence of food. On the other hand, when there are very few bats in an insect-patch, it might be beneficial for these bats to exhibit a patch defense behavior. Indeed, most cases of patch defense described in the bat literature discuss a situation when a single bat deters conspecifics arriving at its patch^14, 45, 46^.

Several previous studies reported that bats change their emission frequencies in response to actual nearby conspecifics^2–5^ or to the playback of interfering signals^6–10^. Researchers have interpreted this behavior as a spectral jamming avoidance response. Explaining all previous studies would require much more than a short discussion. We will thus suggest two alternative hypotheses that could explain these findings and should be further pursued. Except for a few exceptions, the great majority of previous studies reported an upward frequency shift, i.e., bats always elevated their frequency. Such a response could be part of the clutter response that is typical for bats when flying in the vicinity of nearby objects. The function of the clutter response^19^ is to improve localization of nearby objects; in this case other bats. A clutter response is characterized by higher signal frequencies and by additional signal adjustments such as a decrease in signal duration, as several of these studies reported^5^. Some of the previous studies where playback experiments^6–9^, in which additional bats were not present. In these experiments the clutter should not have increased and thus should not have caused a frequency shift. One possibility is that bats approach the playback speaker (as many bats do^47–49^) and thus clutter actually did increase in these experiments. Another possible explanation is based on the Lombard effect, i.e. raising emission intensity in the presence of noise, which is well documented in many mammalian species^44, 50–53^, including bats^2, 13^. It is documented that increasing the signal frequencies could be a by-product of the increase in signal amplitude^54–56^. In both hypotheses, the change in frequency does not aim to decrease spectral overlap and thus cannot be considered a spectral jamming avoidance response. Importantly, the great majority of these studies found an increase in frequency in the presence of conspecifics independently of the frequencies of the two nearby bats. In fact, we also found this in two previous studies^5, 12^. We suggest that such an increase can be explained by the well-documented echolocation response of bats to nearby objects (in this case other bats). We used our simulations to test this hypothesis by reproducing the analysis performed in these previous studies. That is, we analyzed the frequencies used by bats when flying alone and when flying with nearby conspecifics (as was done in previous studies). Indeed, our simulations show that bats’ average frequency in the presence of conspecifics would rise by as much as 400 Hz in comparison to when flying alone, as reported in previous studies, although they are not performing a jamming avoidance response (see Fig S 6). Our results thus explain the findings of most of the previous studies that reported a JAR.

Our work demonstrates the power of simulations to reveal new insight into complex biological systems that are difficult to examine and analyze otherwise. Our model shows both positively and negatively that jamming is less of a problem than previously suggested. On the positive side, it proves that bats can successfully hunt in the presence of other bats without applying any JAR, while on the negative side, it shows that applying a JAR has no significant impact on prey detection or SNR. Similar (modified) simulations can be used in the future to examine many additional fundamental questions in echolocation and to provide insight that may allow us to interpret previous behavioral results and to design better behavioral studies.

## Methods

### General

The MATLAB model simulates the flight and echolocation behavior of *Pipistrellus kuhlii* bats. This small insectivorous bat (approximately 5–9 g) is common in the Mediterranean region and is often observed in groups of ∼5 individuals foraging around a street-light ^13, 15, 24, 57^. Our hunting-ground is a 10×10m^2^ 2D area with no obstacles. Our model consists of three major modules: the prey module, the bat module, and the acoustics module. The prey module controls the flight maneuvers of the simulated moths. The bat module simulates the bat’s behavior and executes the following processes: decision making, echolocation behavior, flight control, and sensory processing. The acoustics module calculates the timing and powers of the received signals from conspecific calls and echoes.

### The Prey Module

The movement of the targets was simulated by a ‘correlated random walk’ model, resembling a flight path of a moth ^21, 22^. The linear velocity has an average of 1m/s and it changes every 100ms according to

Equation 2, where the velocity’s change (Δ_v_) is sampled from a normal distribution: 0±0.1m/s (mean±SD, standard deviation). The velocity is bounded between 0.8 and 1.2m/s. The flight direction is determined according to Equation 3. In this equation,*θ̇*_*n*_, the angular velocity during a 100ms section is a normal random variable with distribution: 0±2 *rad*/s (mean±SD), and τ is the sample time of the model (0.5 ms). The starting position of each moth is drawn from a uniform distribution across the 2D area, its initial flight-direction is random between 0 to 2π radians, and the starting velocity is also normally distributed: 1±0.1m/s (mean±SD).

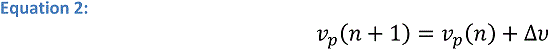

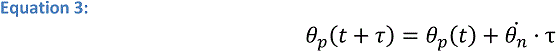

The simulated prey does not detect or respond to the pursuing bats. When a moth reaches the borders of the confined area, it changes its flight angle by *^π^⁄_2_ rad* relative to the border (to return into the foraging area). In order to keep a constant prey-density during the simulation, each time a bat captures a moth, a new moth is added to the environment at a random position with a random flight direction.

### The Bat Module

#### 1) Decision making

The echolocation behavior and flight-control of the simulated bats are illustrated in Fig S 7. At the beginning of a simulation, each bat starts foraging in a random position and transmits echolocation signals with ‘search’ phase parameters (Table 1).

**Table 1:** the echolocation parameters in the different hunting phases

| Flight phase | Search | Approach |  | Buzz |  |  |
| --- | --- | --- | --- | --- | --- | --- |
| Parameter |  | Start | End | Terminal 1 Start | Terminal 1 End | Terminal 2 |
| Inter Pulse Interval [ms] | 100 | 70 | 35 | 18 | 6 | 5 |
| Pulse Duration [ms] | 7 | 5 | 2 | 2 | 1 | 0.5 |
| Terminal Frequency [kHz] | 39 | 39 | 39 | 39 | 39 | 19 |
| Chirp Bandwidth [kHz] | 8 | 35 | 30 | 30 | 20 | 20 |
| Pulse Power [dB-SPL] | 110 | 110 | 90 | 90 | 80 | 80 |

After emitting an echolocation signal, the bat processes all of the sensory inputs (echoes and masking sounds) and decides its next step. The rules of the decision making are as follows: (1) If the bat’s flight-path comes too close to another bat (less than 20 cm) or to the borders of the area, it avoids them and changes its flight direction and velocity. (2) If one or more prey items are detected, the bat chooses the closest one and executes a hunting maneuver. (3) Else, the bat continues searching. According to its decision, the bat adjusts its flight control and echolocation behavior. This decision process is executed every inter-pulse-interval (i.e., between the emissions of two echolocation signals).

#### 2) Echolocation behavior

The echolocating behavior of the simulated bats was modeled based on a rich body of literature^15, 25, 43, 58^. The foraging behavior of insectivorous bats is divided into three main phases: ‘Search’, ‘Approach’ and ‘Buzz’^23^. Each phase is characterized by a different set of echolocation parameters. Our model follows the echolocation and hunting behavior of *Pipistrellus kuhlii* based on field studies^15, 24^ and on lab recordings of eight *P. kuhli* bats trained to search for and land on a static target in a flight room (4.5×5.5×2.5m^3^). The bats in the simulation emit Frequency Modulated (FM) down-sweep signals (mimicking *P. kuhliis’* first harmonic, see Table 1).

Once prey is detected, the hunting phase is defined by the distance to the target. Based on the literature, ‘Search’ switches to ‘Approach’ at 1.2 m from the target and ‘Approach’ switches to Buzz’ at 0.4 m. ‘Terminal Buzz 1’ changes to ‘Terminal Buzz 2’ at 0.2 m. During each behavioral phase, the IPI, pulse duration, bandwidth, and pulse power (in dB) are reduced linearly between the start and end values (Table 1).

### Alternative *JAR models*

Bats emit linear frequency modulated (FM) down-sweeps. We tested three versions of the model: (1) All bats have the same baseline signal, meaning that if two bats are in the same phase and equal distances to targets, their signals will be identical. (2) Each of the simulated bats emits a slightly different echolocation signal; differing by randomly selecting the terminal frequency from a normal distribution, with a mean set to 39 kHz, and a standard deviation of 4 kHz in the ‘Search’ phase. The bandwidth of the signals is constant between bats, so the entire frequency range shifts according to the terminal frequency. This frequency range is in line with the variance of the terminal frequencies reported in the field for this species ^15^. (3) To examine the effect of JAR in the third version of the model, bats used active JAR. They evaluated whether their echoes were jammed (e.g. the masking signal blocks the detection of the prey) by conspecifics’ signals. In such cases, they shifted their terminal frequency upward or downward in steps of 2 kHz, in the opposite direction of the masker (i.e. if the masker’s frequency was lower than their own, they would raise their frequency). These frequency-shifts are in line with the findings of studies reporting evidence of JAR^5, 6^. The bats kept transmitting the modified signal for five consecutive signals, and if during that period another echo was jammed, the bats would shift their frequency again in the proper direction. The terminal frequencies were bounded between 35 and 43 kHz (i.e., there was no shift beyond these boundaries).

#### 3) Flight Control

Before a prey is detected, simulated bats fly according to a ‘correlated random walk’ path, with a constant linear velocity 3.5m/s and a random change of direction, sampled from a normal distribution of angular velocities: 0±1 rad/sec (mean±SD)^27, 32, 58^. A new angular velocity is sampled before each echolocation emission and the bat turns according to this velocity until the next emission. Once a target is detected the bat turns toward the prey by changing its angular velocity according to its relative direction to the target, using a delayed linear adaptive law described in ^27, 59^. This dependency is described in Equation 4, where *θ̇*_*bat*_(t + τ) is the angular velocity of the bat in the next time-sample (i.e., time t + τ), k_r_ is a gain coefficient, limited by the maximal acceleration of the bat (set to 4m/s^2^), and *Ø_target_(t)* is the current angle between the target and the bat.

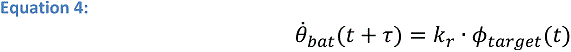

We neglect head movements; the original model^20, 27, 59^ refers to the gaze angle (i.e., the angle between the head’s direction and the target), but for simplicity, we assume that the head and body are aligned. Even though we know the head and body are not always aligned; this assumption does not affect the behaviors tested in this study. To keep its direction aligned with the target, the bat typically slows down when the angle to the target (Ø_target_) is large, and accelerates linearly when the target is straight ahead ^18, 60^. To model this, we implemented a velocity-model suggested by Vanderelst and Peremans to simulate this behavior ^27^, described in Equation 5. V_phase_ is the maximal velocity in each behavioral phase (3.5m/s in the approach and search phases, and 2m/s during the buzz phase). Like the direction, the bat adjusts its velocity after each inter-pulse-interval. Indeed, the accelerations and turning rates of the simulated bats correspond well to those reported in the field ^18, 60^, see Fig S 1. A successful hunt (capture) is achieved when the bat is less than 5cm from the prey^27^.

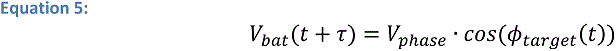

#### 4) Sensory Processing

The simulated bat detects and estimates the range and direction of objects in the environment (prey and obstacles), based on the incoming acoustic input. Echoes will only be processed if they cross the auditory threshold set to 0 dB-SPL based on the literature ^42, 61–63^ (we also tested the influence of that threshold between 0-30 dB-SPL, see results, Figure 5C). We define such detected echoes as ‘Pre-Masking Echoes’. Next, we calculate the effect of masking using two different detection-models: the **correlation-receiver** which is well-well studied theoretical reference model, and the gammatone **filter-bank receiver** which represents the temporal reaction of the inner ear to auditory signals. After the preliminary detection, the bat chooses its target again, from the non-jammed echoes.

The **correlation-receiver** is based on a similarity between the bat’s own transmitted signals and the received signals^30, 64^. The detector calculates two correlations: (i) the self-correlation between the echo and its own transmitted signal. (ii) The cross-correlation between the masking interference signal and its own transmitted signal. For the echo to be detected, the self-correlation peak should be higher than the cross-correlation peak with more than the ‘forward detection threshold’ (set to 5dB) if the cross-correlation peak is within 3ms before the echo, and higher than the masking peak by more than the ‘backward detection threshold’ (e.g. 0db), if the masking signal arrives within 1ms after the wanted signal (see Fig S 8). The periods and thresholds were defined according to ‘the law of first wave-front’^63^ ch. 2.4.5, ^65^ch. 3.1 and ^66^, and comprise lower boundaries of real bats’ abilities to cope with masking. For simplicity, we used constant thresholds within each window. Even if the echo is detected, masking signals may still degrade the accuracy of the sensory estimations, i.e., the distance and angle, see below. Masking sounds arriving outside the reception window do not interfere with detection.

The **filter-bank receiver** is based on the bat hearing model described in Weißenbacher and Wiegrebe (2003)^39^. The detector consists of an 80-channel gammatone filter-bank with frequency bandwidths simulating the tuning curve of the inner-ear^36, 67^. The impulse response of each channel in this model is given by Equation 7; where *n* is the filter order (set to 4), *b* is the time constant of the impulse-response (set to 0.15f_c_), f_c_ is the center frequency of the channel (Equation 7). The signal is filtered by a lowpass filter, which keeps only the envelope of the signal (‘envelope detector’).

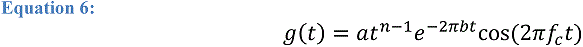

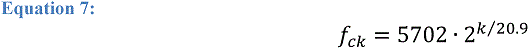

Then, we shift each channel in time to compensate for the delay time between the emission time and the response time of each channel, according to the chirp’s slope. Finally, we sum the time-compensated filter-channels and look for peaks in the integrated signal. We define prominent peaks as those that are higher than their surroundings by more than the acoustic noise level. All prominent peaks that are also higher than the detection-threshold, set to 0 dB-SPL, are referred to as potentially detected echoes, and their distance from the bat is estimated relative to the peak detection-time, see Fig S 9. Note that these peaks might be a result of desired echoes or masking signals and their amplitude will be determined simply by running them through the filter-bank model.

Like with the correlation-receiver, for each transmitted signal, we implemented the filter-bank receiver twice: (a) only on the echoes from the bat’s own emitted signal, and (b) on both masking signals and echoes. We compared (a) and (b) detected peaks as described above and then defined the following criteria: a jammed signal is a peak that was detected in (a) but not in (b); the time-estimation error is the difference between the estimated peak-time in (b) and the actual received time of the reflected echo; and false-alarm occurs when a bat decides to pursue a ‘target’ that was detected only in (b), and therefore is not a prey echo, but a masking signal.

Note that ‘correlation-receiver’ assumes that the bat can differ between desired prey echoes and masking signals and echoes from conspecifics ^13, 68^. On the other hand, the ‘filter-bank receiver’, only assumes that bat estimates the times of its own transmitted signal in each channel.

The SNR (Signal to Noise and interference Ratio) is calculated for each detector by Equation 8.

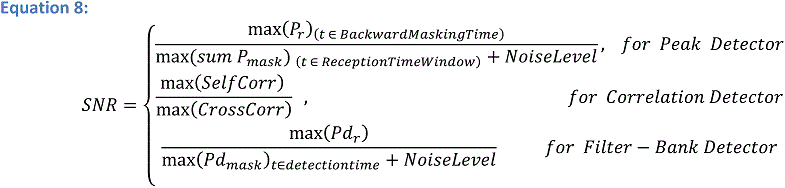

To analyze the effect of signal duration on performance, we modified the model by implementing the correlation-detector in the detection process too, before the masking calculations. Here, the received echo was first correlated with the emitted signal and then it was compared to a detection-threshold. The detection threshold for the correlation was set to 15 dB-SPL, which equals to the maximum of the autocorrelation of an 8ms ‘search’ signal.

After the detection process, the bat estimates the range and the Direction Of Arrival (DOA) of the reflecting object, based on all of the received signals (echoes and masking signals). The range estimation is based on the acoustic two-way time-travel of the signal with an error (Equation 9), comprised of two elements: the bat’s accuracy in measuring time, and a noise term which reduces with increasing SINR (Equation 8, calculated using either the Correlation or the Filter-bank model). For simplicity, because all bat signals are FM-chirps, we use an error that is independent of the signal’s parameters: the term *k_1_/_SINR_*, where *k_1_* is a coefficient set to scale the error to values of ±1cm at SNR of 10dB, based on behavioral studies^30, 63, 69, 70^.

The independency of the signal’s parameters is reasonable because all bat signals in our simulation were similar. The term *Time_Res_* (in Equation 9) is sampled from a Gaussian distribution with a mean ± SD of 0±50 microseconds, equivalent to a range error of 0.85 cm, which is a low boundary estimation of bats capabilities to measure distance^54^. ‘c’ is the speed of sound: 343m/s.

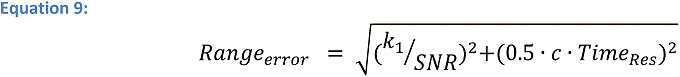

The estimation error of the DOA, see Equation 10, includes an error which depends on the DOA (*k_3_ + k_4_ · sin(Ø)* in Equation 4); set such that the error equals 1.5° at 0° DOA, and error of 10° at 90° DOA, and a Gaussian noise term: 0°±1° (mean ± SD) at SNR=10dB, see *k^2^/_SINR_* in (“Hearing by Bats”, Chapter 3.1, ^63^) ^71^. Fig S 10 depicts the histogram of the consequential range errors and DOA errors.

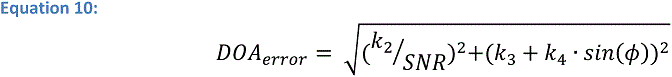

In general, our model intentionally underestimates the bats’ actual performance, and thus real bats are likely to cope with acoustic interference even better than our simulated bats: (1) we simulated monaural bats, while real bats use two ears with spatial selectivity^31^. (2) We assumed a low detection threshold (0 dB-SPL), thus the bats were more susceptible to masking (see Figure 5 C2). (3) We chose long backward and forward masking windows (3ms, 1ms), and low jamming thresholds: 0dB for backward masking, 5dB for forward masking (e.g. a masking signal that was received first, even if it is 5dB lower than the desired echo, will totally jam it). (4) The model implied a pulse-by-pulse detection and estimation strategy with no memory, therefore, temporal miss-detections caused by momentary jamming had a very substantial effect on hunting attempts.

### Acoustics Calculations

The estimated intensities of the reflected echoes based on the sonar/radar equation (^72^, pp. 196–198), shown in Equation 11, angles and distances are defined according to Fig S 11.

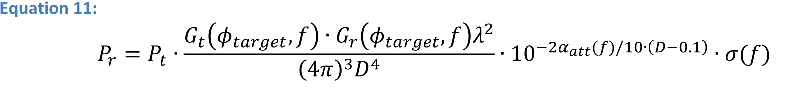

Where, P_r_: Power (intensity) of the received signal [SPL]

P_t_: Power (intensity) of the transmitted call [SPL]

Ø_target_: Angle between the bat direction and the prey [rad]

G_t_(Ø, f): Gain of the transmitter (mouth of the bat), as a function of azimuth and frequency (*f*)

G_r_(Ø, f): Gain of the receiver (ears of the bat)

D: Distance between the bat and the object [m]

α_att_(f): Atmospheric absorption coefficient for sound [dB/m]

σ(f): Sonar cross-section of the target [m^2^]

λ: The wavelength of the signal [m]

The transmitted power (P_t_) of a search signal equals 110 dB-SPL at a distance of 0.1 meters from the source ^71^. P_t_ during the hunt varies according to the distance to the prey (see Table 1). The transmission gain of the bat’s mouth Gt(ϕ, f) is modeled by a circular piston source ^73–75^, given in Equation 12. The directivity of the emitted call depends on the ratio between the wavelength of the signal (λ) and the radius of the emitter (mouth), ‘a’ (set to 3 mm ^74^); J_1_ is the Bessel function of the first order and k= 2π/λ. G_0_ of the mouth (at the head direction) is set to 1, matching the measurements of intensities of bat’s calls from a distance of 0.1m.

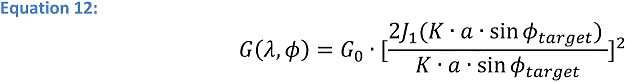

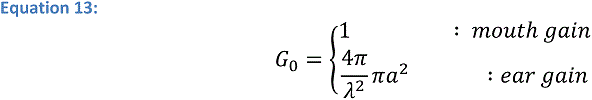

To estimate the gain of the ears (Gr(Ø) in Equation 11), we modeled them as circular planes, using the piston model ^58^, with ‘a’ (the ear radius) set to 7 mm, matching *Pipistrellus kuhlii*’s ear size ^76^. G_0_, the maximum gain of the ear, is defined by Equation 13 (^72^ Equation 5.36, pp. 181), where A is the geometric area of the ear. Since *Pipistrellus* bats do not move their ears, this is a reasonable estimate.

The original piston model is symmetric: the back-lobe is equal to the front-lobe. In our model, bats receive signals from the back hemisphere too, therefore we modified the piston model for the back hemisphere (for transmission and reception), and it is shaped by the piston model with additional attenuation of 0-20dB, increasing linearly (in dB) from ±90° to ±180°, relative to the bat. The modified piston model for ear and mouth are illustrated in Fig S 12.

α_att_ in the sonar equation (Equation 11) is the atmospheric absorption coefficient, set to temperature 20°C and humidity 50%, is approximated by Equation 14^58^.

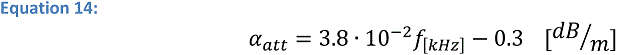

σ, the Radar (sonar) Cross-Section (RCS, or ‘target strength’) of the moth, is modeled as a disc with a radius (‘r’) equals 2 cm, equally reflecting in all directions. We apply the approximation of the RCS for this type of reflector^77^, given in Equation 15, where A is the geometric area of the disc (A=πr^2^). ‘r’ was set to 2cm, simulating a moth’s wing-length. This approximation is in line with measurements of the target strength of medium-sized insects ^42^[figure 1h], ^43^.

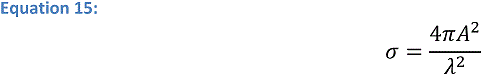

The receiving time-delays of the echoes are the 2-way duration of the signals propagating from the emitter to the target, see Equation 16, where, ‘c’ denotes sound velocity, D is the distance between bat and target, and t is the overall travel time from the emission to the reception.

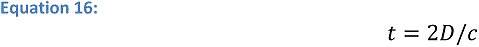

The power of masking signals, i.e. signals transmitted by conspecific bats (t_x_), received by the bat (r_x_), is calculated by Equation 17, where *P_mask_* is the emitted power of the masking signal, and the angles are depicted in Fig S 11.

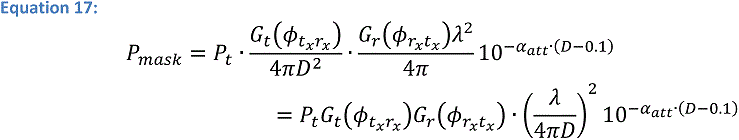

In addition to its own echoes (Equation 11) and conspecific signals (Equation 17), each bat also receives echoes that are reflections of objects from conspecific signals. The intensities of these potentially masking sounds are estimated by modifying Equation 11 (sonar equation), with the suitable angles and distances. This calculation is described in Equation 18, and the angles are defined in Fig S 11, arrows 3-4.

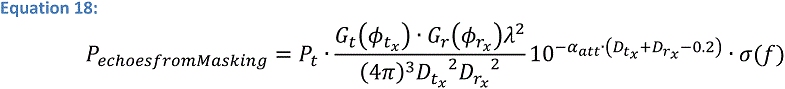

### Data Analysis and Statistics

In each scenario that we simulated, we tested the effect of the varied parameters on hunting performance using ANCOVA, ANOVA or multiple regression, depending on the type of the parameters (e.g. continuous or categorical) and their number. The Statistics were calculated using Standard Least Squares method with JMP 14 and MATLAB R2018b.

For testing the significance of the masking effect, we used a different procedure. Because the masking effect is a ratio (see Equation 1) we had to compute its SD. Thus, for each set of conditions, we estimated the standard deviation of the masking effect, using Equation 19 ^78^. In order to determine whether the masking effect significantly dependent on the tested parameters, we simulated 100 points in each scenario with the average and SD calculated above, and executed ANCOVA to estimate the F test, for the simulated scenario. We repeated this process 1000 times in each scenario and used the average results of the F tests and p-values as the statistics. This process of repetitions was executed only for the statistics of the masking effect.

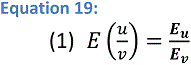

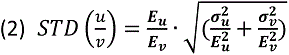

We also evaluated the ‘Jamming Probability’ as the proportion of pursued prey echoes (echoes of prey the bat chose to pursue) that are blocked by masking signals (Equation 20). Note, that this ratio is not the proportion of all the jammed echoes to all detected echoes, because for each signal there can be several detected echoes from different prey items, but only one (maximum) is pursued.

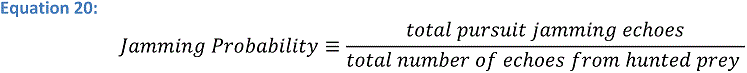

## Supplementary

**Fig S 1:**
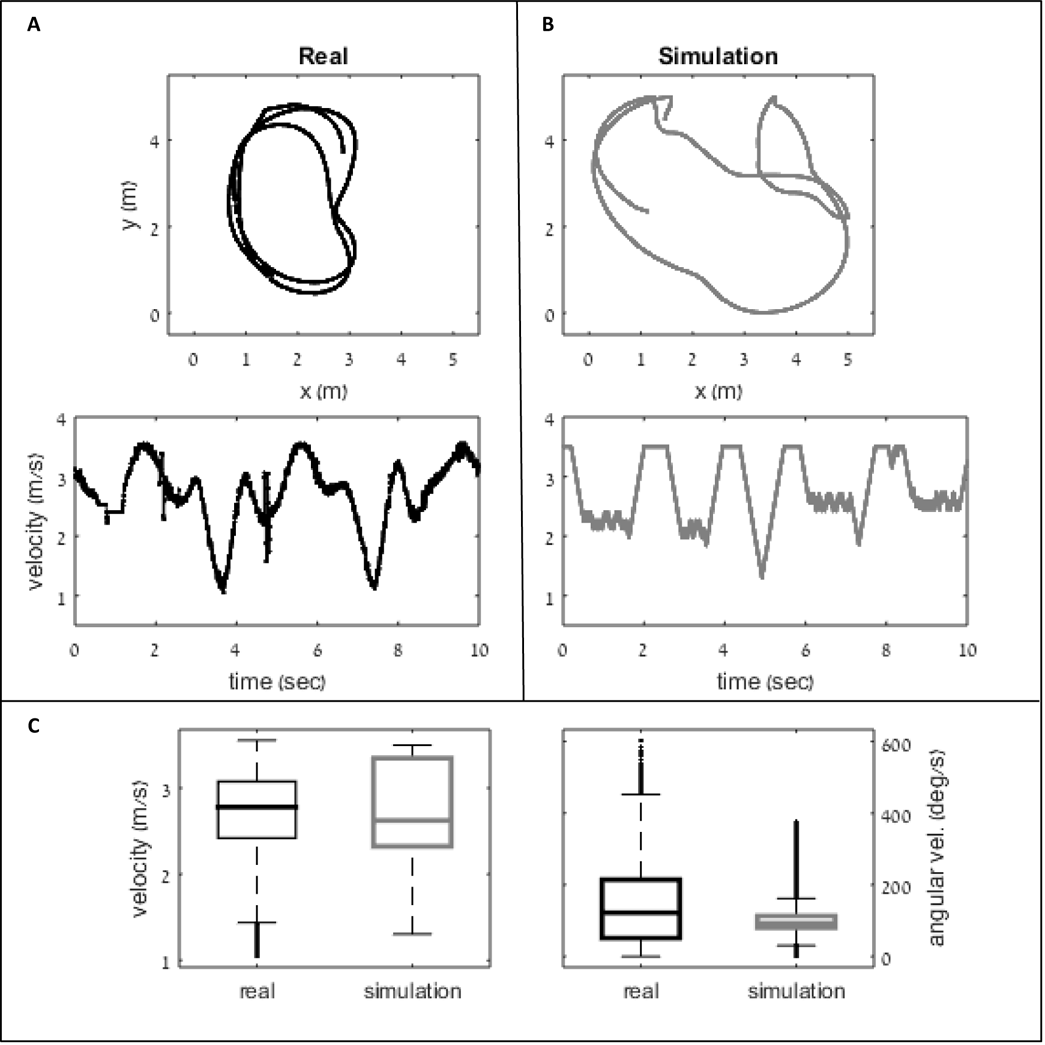
Simulated and real bats flight characteristics. (A) A typical example of one real bat flying with two conspecifics in a closed flight room (5.5×4.5×2.5 m^3^) with a feeding platform that they were trained to land on. The upper panel illustrates the flight trajectory (two dimensional) of the bat in 10 seconds. The lower panel illustrates the linear velocity of the bat as a function of time. **(B)** The flight-path and velocity of one simulated bat flying in a group of three bats with a static target in a 5×5m 2D arena. **(C)** Boxplots (median, q1, q3 and outliers) of the linear-velocities and angular-velocities of the real bat (black) and simulated bat (gray). Note that both linear velocity and angular velocities of the model are in the range of natural velocities in similar conditions. Additionally, the acceleration of the model was bounded to 4m/s^2^ while real bats’ acceleration was below 5m/s^2^ during 95% of the time.

**Fig S 2:**
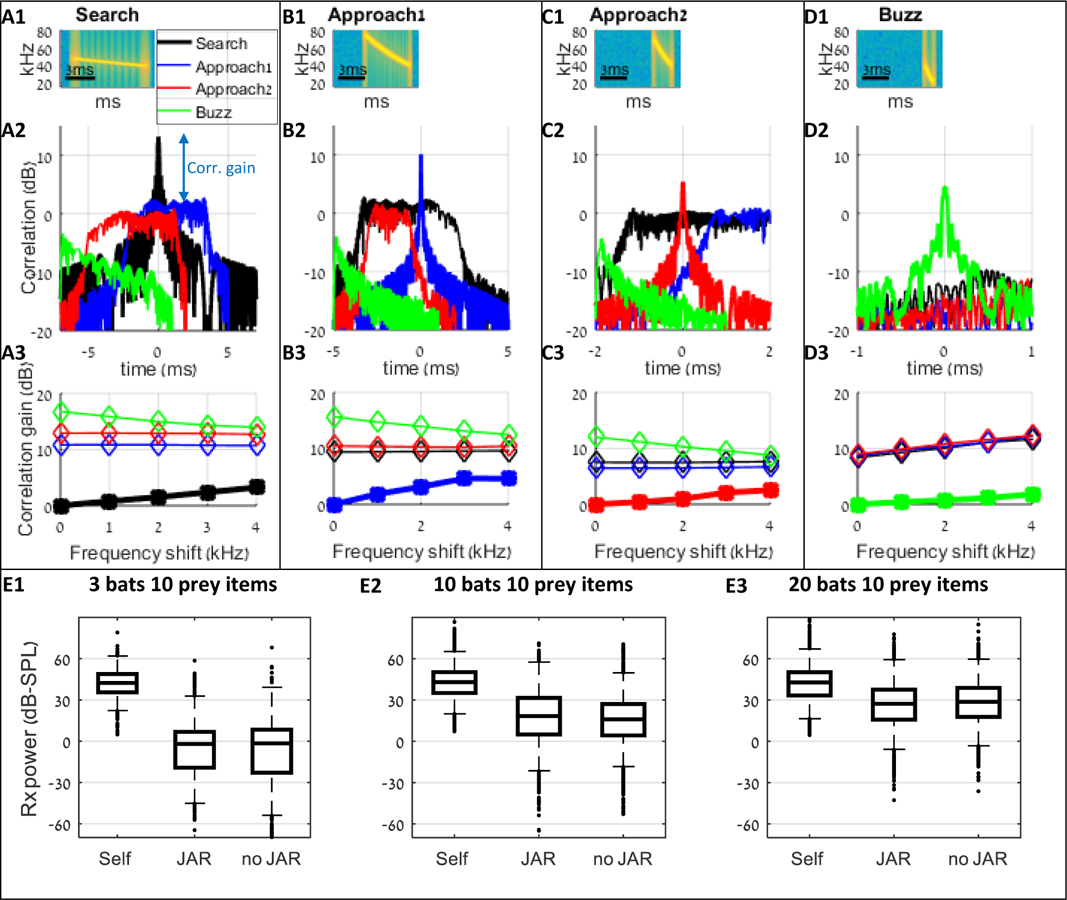
Analysis of correlation gains between signals of search (A), approach (B-C) and buzz. (D) phases. The effect of JAR behavior can be quantified by examining the similarity between the desired prey echoes and the masking signals. During the search phase, the correlation between the desired echo and the emitted signal is on average ∼10dB higher than the correlation between the emitted signal and masking signals produced by bats in other behavioral phases. Additionally, there is no benefit in shifting the frequencies of the emitted signal to cope with masking from those signals. If the masking signal is a search signal (as the emitted one), shifting the frequency by 2kHz has a limited improvement of 2dB at most (A3-D3). As a result of this constant spectral variability, applying a spectral JAR to shift the frequencies of the emitted signal does not affect the correlation with the masking signals (E1-E3). **Panel A1** depicts the spectrogram of a transmitted search signal with a duration of 7ms and bandwidth of 8kHz (Table 1). **Panel A2** illustrates the correlation function between the transmitted signal and potential masking signals of bats at different behavioral phases (black - the correlation with another search signal, blue and red-with two different approach signals, and green - with a buzz signal). All signals were received with the same power. The correlation gain is defined as the difference between the peak of the auto-correlation and the maximum of each cross-correlation function (blue arrow– the correlation gain for the search signal relative to the approach1 signal). The correlation gain of the transmitted signal with a search signal is 0dB by definition. To examine the effect of a JAR response, we gradually shifted the terminal frequencies of the masking signals and calculated the correlation gain for each frequency shift. **Panel A3** describes the correlation gain as a function of the frequency shift. The gain improvement due to a JAR response for signals from the same phase (black –‘search’) is ca. 2dB per 2 kHz shift; there is no increase in the correlation gain for signals at other phases. **(B-D)** Same as **‘A’** for additional signals: (**B**) approach1 (‘the first approach signal’ with duration of 3ms and bandwidth of 35kHz), **(C)** approach2 (duration: 2ms, bandwidth: 3kHz), and **(D)** buzz signals (duration: 1ms, bandwidth: 20kHz). **(E) Panels E1-E3** present the results from actual simulations of the received signals after correlation with the emitted signals. Three conditions are shown: ‘Self’ depicts the correlation of the echoes from the target and the transmitted signal. ‘JAR’ depicts the cross-correlation of the received masking signals, when bats applied spectral JAR. ‘No JAR’ is the correlation with the masking signals without JAR. Spectral jamming avoidance did not significantly reduced the masking signals in comparison to the no-JAR scenarios (permutation test, effect-size=-5.8dB /-1.8dB / 0.7dB, p = 0.03*/ 0.49/ 0.41, for 3, 10, 20 bats in 100m^2^, sample-size=150, 1000 permutations, 100 repetitions). Negative effect-size means that the no-JAR bats had a better correlation than the JAR bats; an asterisk (*) indicates a significant difference (P<0.05).

**Fig S 3:**
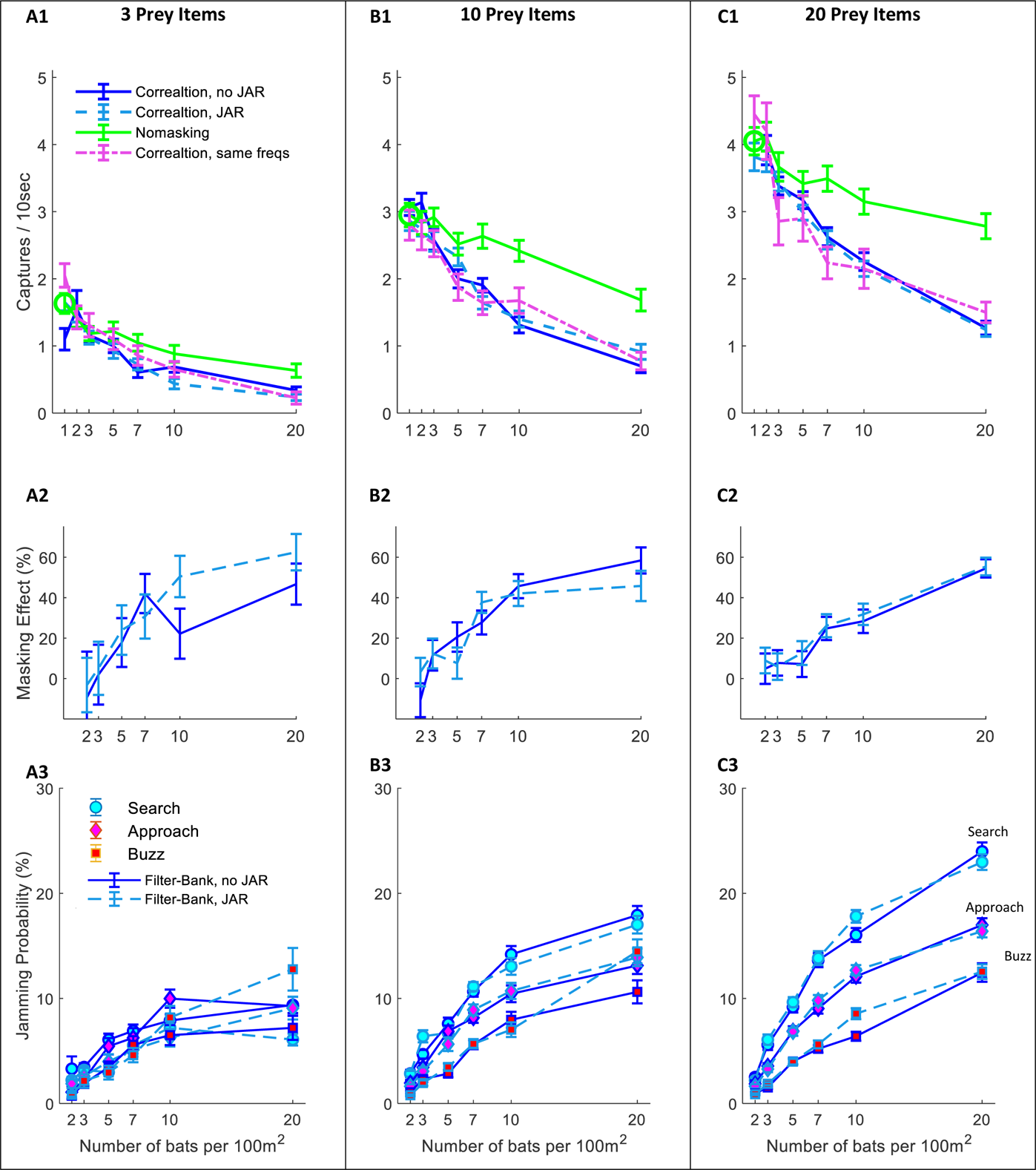
Hunting performance for the filter-bank receiver. The hunting success rate is presented for three prey densities per 100m^2^: **(A)** 3 prey items; **(B)** 10 prey items; **(C)** 20 prey items. Line colors and styles depict the performance of different receiver models: green-no-masking; solid blue – filter-bank receiver with random signal variability; dashed turquoise - filter-bank receiver with JAR response; dash-dotted magenta– filter-bank receiver without signal variability. **Panels (A1-C1)** depict the performance as a function of bat density; a green circle shows the performance of a single bat. In the no-masking condition (green lines) there is a significant degradation in performance as the bat’s density rises from 1, and it reaches to maxima of ca.: 60%, 43%, 32% where, for 20 bats and 3,10,20 prey items per 100m2, respectively. There was no significant difference in performance when applying or not applying spectral JAR-compare turquoise and blue lines (One-way ANOVA: F2,1007=0.467, p=0.529; F2,1355=0.651, p=0.42; F2,1529=0.207, p=0.645. **Panels (A2-C2)** show the masking-effect on hunting, i.e., the relative decrease in hunting relative to the ‘no-masking’ condition. The masking effect for 10 and 20 bats per 100m2 was between 40%, and 60%, respectively. **(A3-C3)** The probability of jamming at different behavioral phases: search (turquoise marker), approach (magenta marker) and buzz (red markers). For prey densities of 10 and 20 prey items per 100m2, there was no significant difference between JAR and no JAR scenarios, in all flight-phases (One-way ANOVA). In scenarios including low prey density (i.e., 3 prey items per 100m^2^), the probability of jamming was low, with a maximum of ca. 10%.

**Fig S 4:**
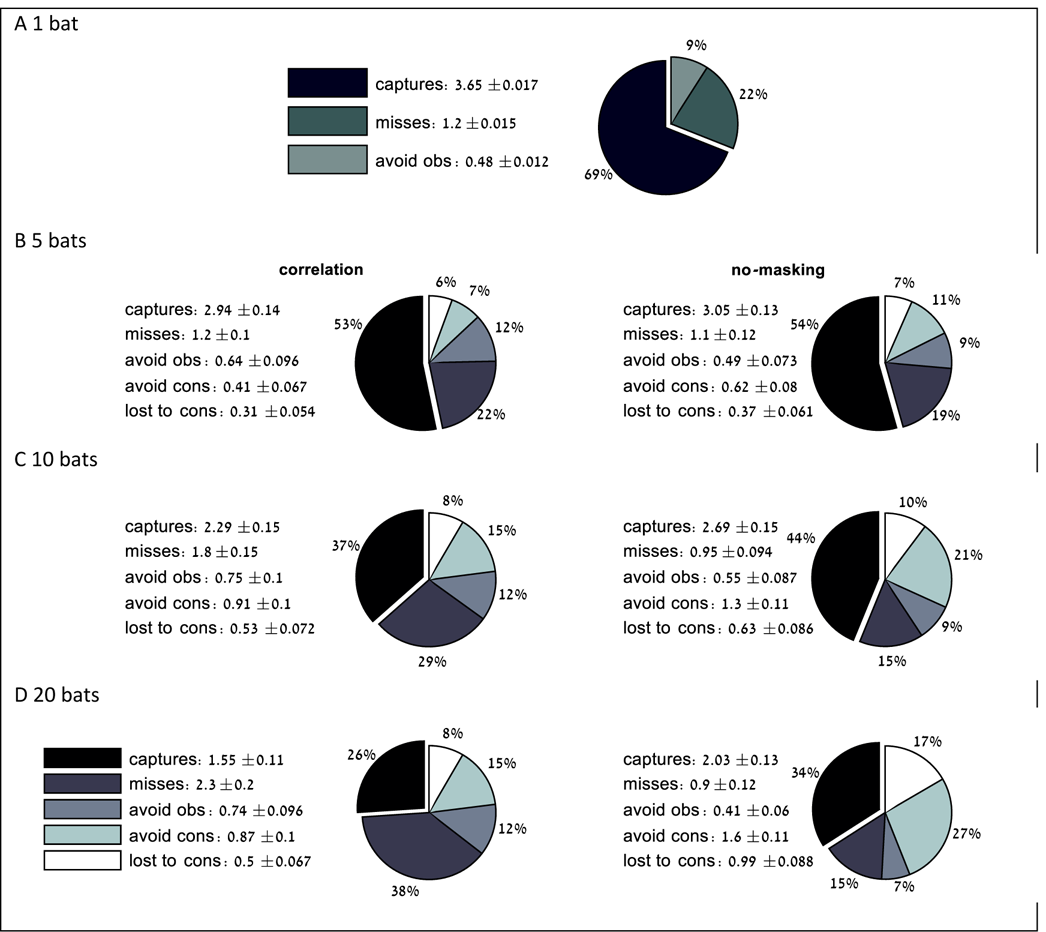
The causes of hunting failures. (A-D) Pie charts present the proportions of successful and unsuccessful attacks with different failure reasons for groups of 1 to 20 bats, with masking (‘correlation-detector’) and without masking. Each chart is divided into the following categories: (i) **captures** - maneuvers that ended with a successful capture; (ii) **misses** - maneuvers in which the bats failed to capture the target because of insufficient maneuvering and/or masking; (iii) **avoid obs** (obstacles) – flight paths that came too close to the borders of the arena and the bat seized hunting; (iv) **avoid cons** (conspecifics) – hunting maneuvers that were ended in order to avoid collision with another bat; (v) **lost to cons** –hunting attempts that were terminated because the prey was caught by a conspecific. The pie charts present the mean and the standard-errors for 100 simulated bats. Each panel includes 100 simulated bats. The absolute rates for each category are presented near the pie chart. In all bat densities the proportion of missing the prey increased when adding sensory masking (One-way ANOVA: effect-size=3%, F_1, 198_=3.21, p=0.075; effect-size=11%, F_1, 198_=20.7, p*<0.0001; effect-size=23%, F_1, 198_=68.8, p*<0.0001, for 5, 10 and 20 bats per 100m^2^). Asterisk (*) indicates significant difference. The permutation test was executed as follows: (1) we randomly sampled 150 points from each data-set (i.e., the received power of ‘JAR’ and ‘no JAR’). (2) We defined the sampled effect-size as the difference between the medians of the samples. (3) We randomly distributed the samples between two groups and calculated the effect-size between the two groups. (4) We repeated step 3 for 10,000 times. (5) The p-value of this sample was the proportion of times the effect-size of the permutations was higher than the sampled effect size (in absolute values). (6) We repeated steps 1-5, for 100 times. The p-value is the median of p-values from all the random samples.

**Fig S 5:**
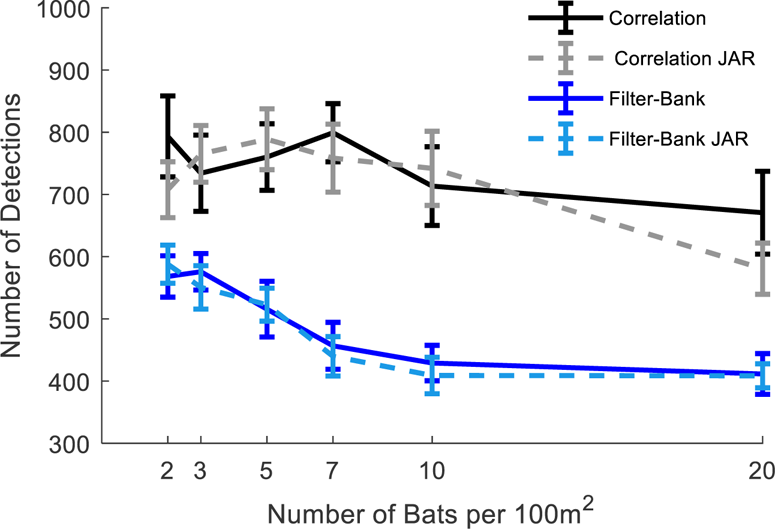
Number of prey detection events. The figure depicts the total number of prey detections per 10 seconds for the correlation-receiver with no-JAR (solid black), the correlation-receiver with JAR (dashed gray), the filter-bank receiver with no JAR (solid blue) and the filter-bank receiver with JAR (dashed turquoise). A JAR had no significant effect on both correlation-receiver and filter-bank receiver: One Way ANOVA: F_1,237_ =0.46, p=0.50; F_1,237_=0.11, p=0.74, respectively. The number of detections with the correlation-receiver is significantly higher than with a filter-bank receiver: One Way ANOVA: F_1,481_=205, p<1e-10.

**Fig S 6:**
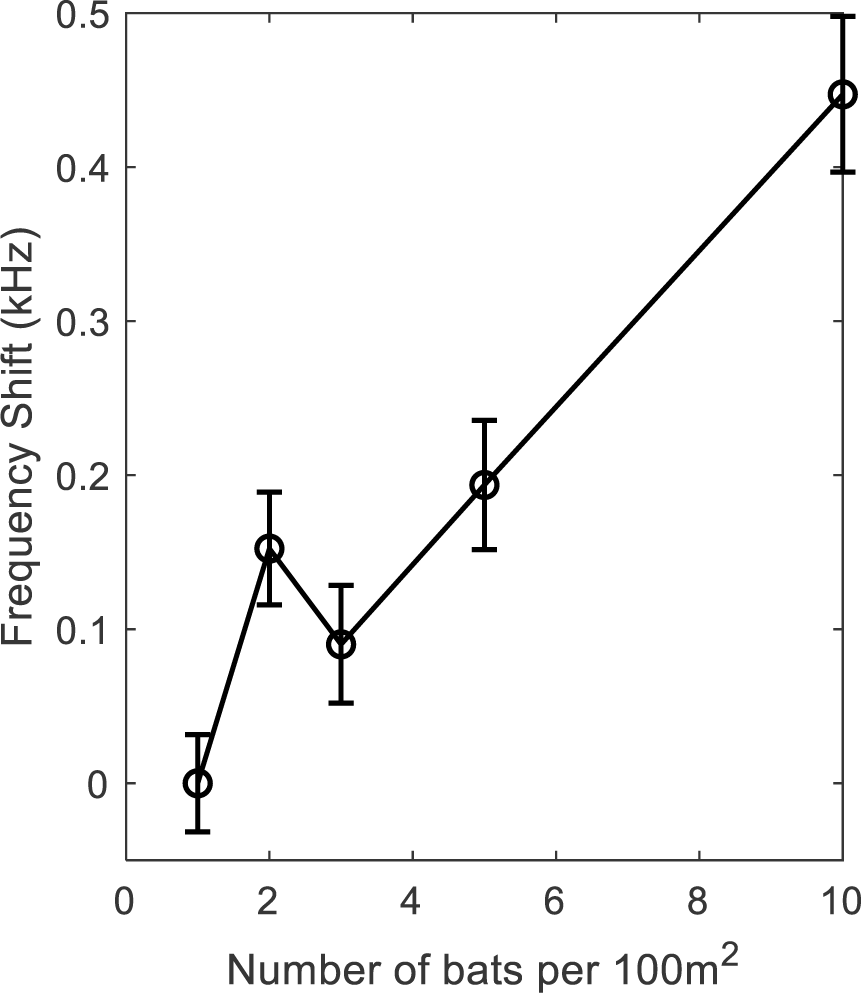
Frequency-shift with no JAR. The shift in the average frequency of the emitted signals, as a function of bat density. All simulations were performed with the correlation-detector, with no JAR. The bats react to conspecifics flying nearby as objects and modify their echolocation behavior according to the distance between them. The results clearly demonstrate a significant rise in frequencies as the density increases (Regression test, F_1,993_=64.4, p<0.0001), there is also a significant rise in frequency when increasing the bat density from 1 to 2 bats per 100m^2^ (t-test: t= 3.15, p=0.0017, DF = 389). All trials with one prey item in the arena. The graph indicates mean and standard errors.

**Fig S 7:**
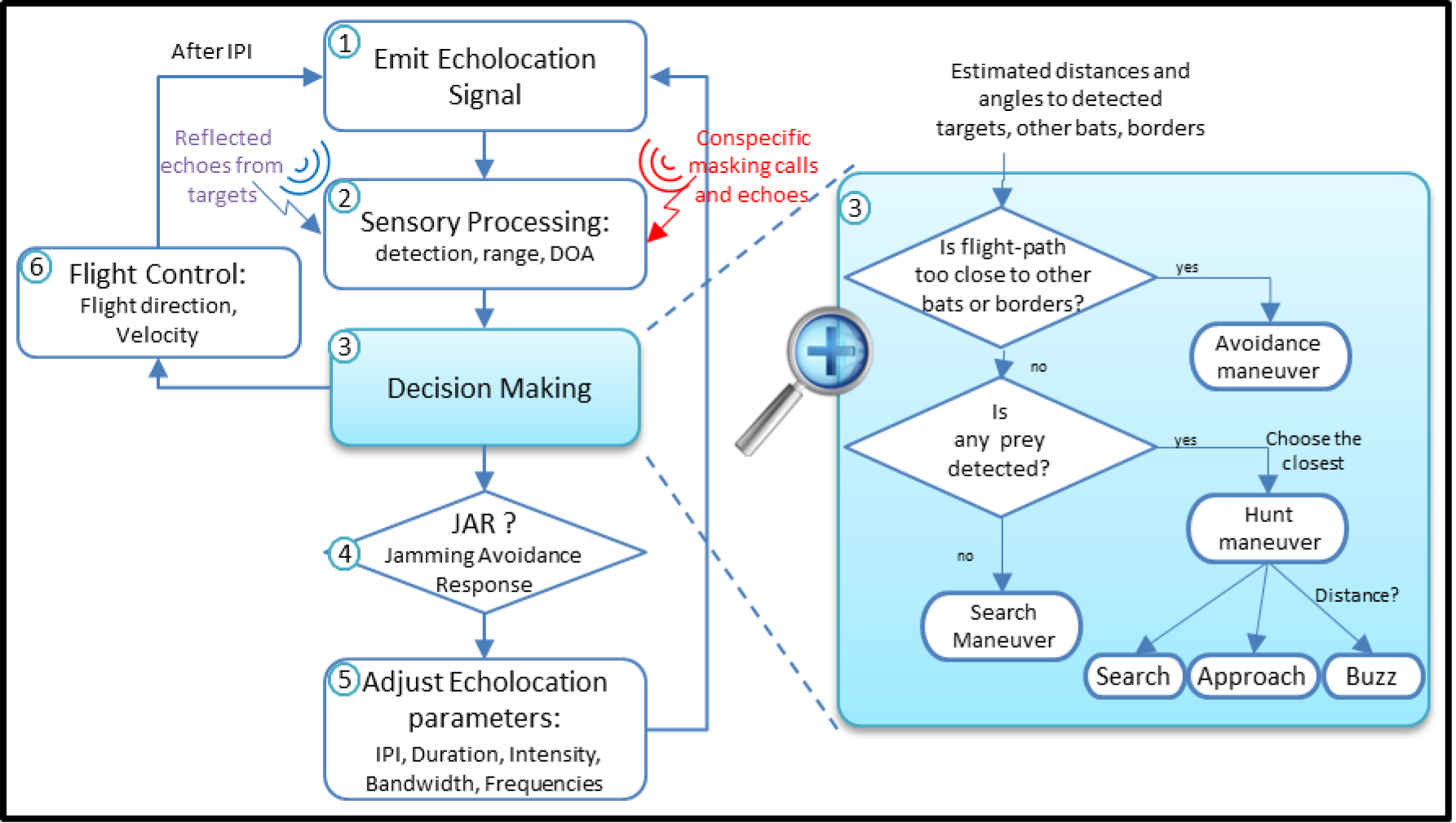
The Bat Module streamlines. (1) The bat emits an echolocation signal. **(2)** Based on the received signals (echoes from objects and masking signals from conspecifics) the bat detects the targets and estimates their relative locations. **(3)** According to the sensory output, it decides its next maneuver: keep searching, pursue prey or avoid obstacles, and determines its next phase: Search, Approach or Buzz. **(4**) If the bat experiences masking and if a JAR is applied, it modifies its echolocation signals. Depending on its decision, the bat adjusts the echolocation parameters **(5)**, and its flight speed and direction **(6)**, and after an inter-pulse-interval, emits another echolocation signal **(1).**

**Fig S 8:**
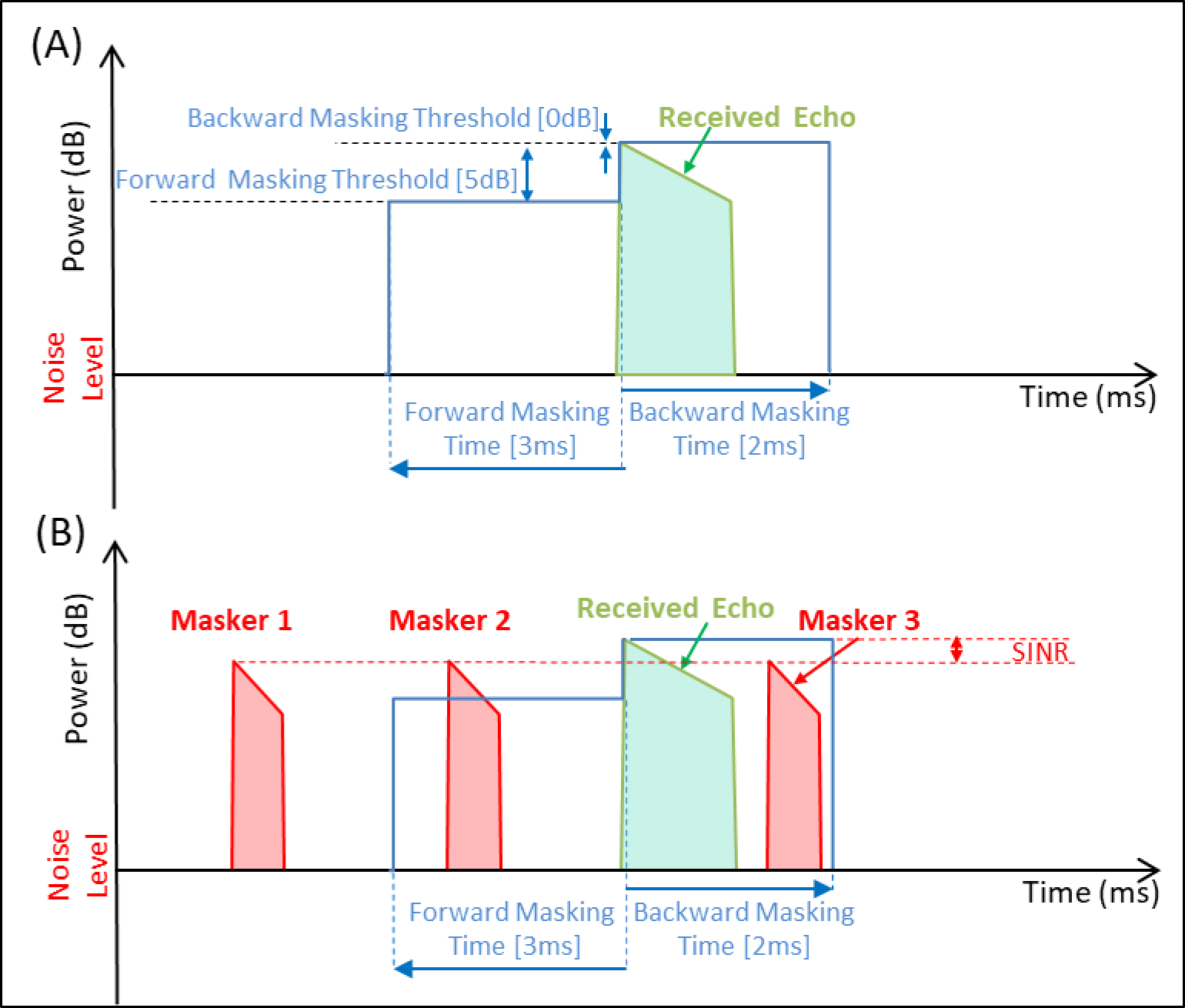
Forward and Backward Masking Criteria. (A) An echo from a target (green) is received by the bat. The reception window (blue) is composed of two periods: the ‘forward masking time’ starts 3ms before the arrival time of the signal, and the ‘backward masking time’ starts at the arrival time and lasts 2ms. The ‘forward masking threshold’ is set to 5dB below the maximum of the received signal during the ‘reception window’, and the ‘backward masking threshold’ is equal to the maximum power of the signal. Any signal (even below the two thresholds) that arrives within the reception window will mask the target’s echo and will reduce its localization accuracy, as a function of the SINR. If a masking signal’s power is above one of the two relevant thresholds, the echo is jammed, and the bat does not detect the target. Panel **(B)** illustrates an example of a received echo with three potentially masking signals (red). Masker 1 does not affect the detection, because it is outside the ‘reception window’. Masker 2 is above the forward masking threshold; thus the echo is jammed and the prey is not detected at all. Masker 3 is received within the reception window, but is below the backward threshold, and as a consequence, the signal is detected but the localization error increases, according to the SINR (Equation 9, Equation 10). Both models of receivers (‘correlation-detector’ and ‘peak-detector’) apply the same ‘reception window, and thresholds. However, the peak-detector uses the maximum power of the received signals to set the thresholds, whereas the correlation-detector uses the maximum of the correlations between the received and the transmitted signals.

**Fig S 9:**
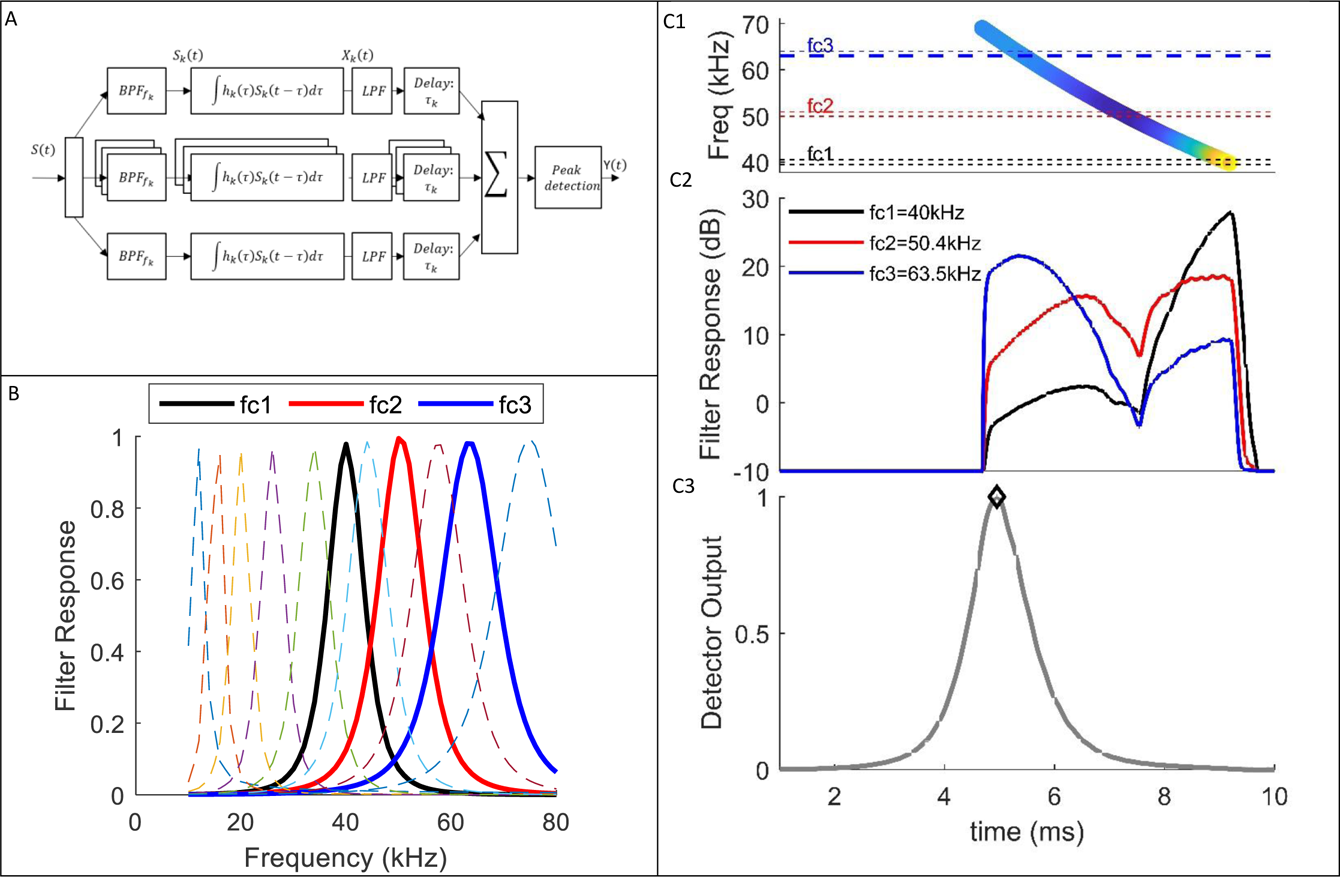
Filter-Bank Receiver. (A) Receiver Scheme: the received signal, s(t), is split to n-channels of gammatone filters. In each channel, the signal is filtered by a band-pass filter, with center frequency fc_k_, and input response h_k_.. The signal goes through a low-pass filter (‘envelope detector’) that removes phase information. Then, each signal is shifted in time (τ_k_) to compensate for the chirp slope (i.e. τ_k_ is equal to the time difference between the start of the transmitted chirp, and the response in each channel). The channels are integrated, and the detector looks for local maxima (i.e. peaks). The receiver estimates the intensities and delays of the detected peaks’ that cross the detection threshold (see text). **Panel (B)** depicts the frequency responses of several gammatone filters (11 out of 80). τ_k_ illustrates the signals in several points at the process: **(C1)** Spectrogram of a reflected echo from prey (S(t)); **(C2)** The response of three gammatone filters (fc1,fc2,fc3), before the time shift; **(C3)** The receiver’s output (Y(t)) after the integration and peak-detection processes.

**Fig S 10:**
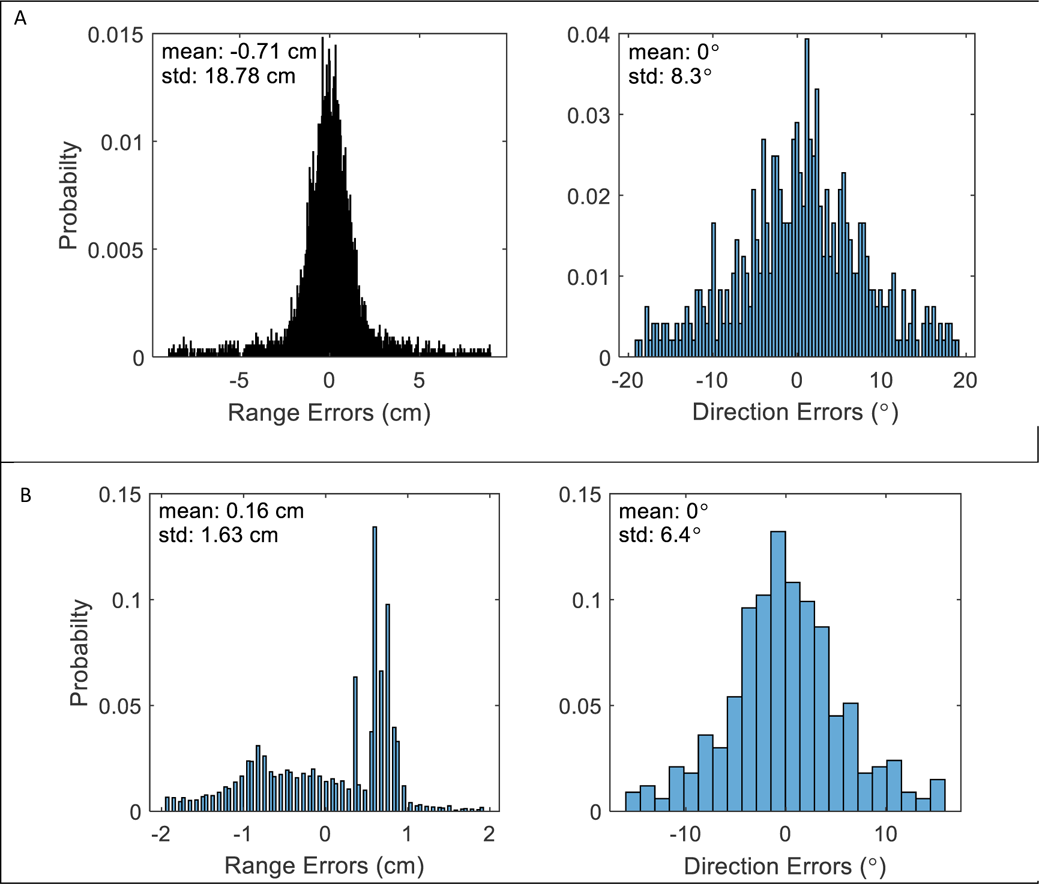
Ranging errors of the Correlation and Filter-Bank receivers. Errors are defined as the difference between the estimated and the actual location (range and direction) of all detected prey, for trials with 5 bats and 10 prey items**. (A) Correlation-receiver:** the range errors (left panel) and direction errors (right panel) are calculated according to Equation 9: and Equation 10:, respectively. The average of the absolute errors is 10.4cm and 9°. More distant objects have lower SNR’s, and, hence, higher estimation errors. The standard error of range errors for detections with high SNR of 10dB is ca. 1cm and the standard error of direction is ca. 2°. **(B) Filter-bank receiver:** the range errors (left panel) are derived from the timing estimations of the peaks of the detected signals (see Methods). Direction errors right panel) are calculated by Equation 10:, as for the correlation-receiver. The average of the absolute errors is 1.1cm, and 5.2^0^, respectively. The error is lower in comparison to the correlation-receiver because the filter-bank receiver only detects higher SNR objects. Apparently, this difference has little effect on the hunting rate.

**Fig S 11:**
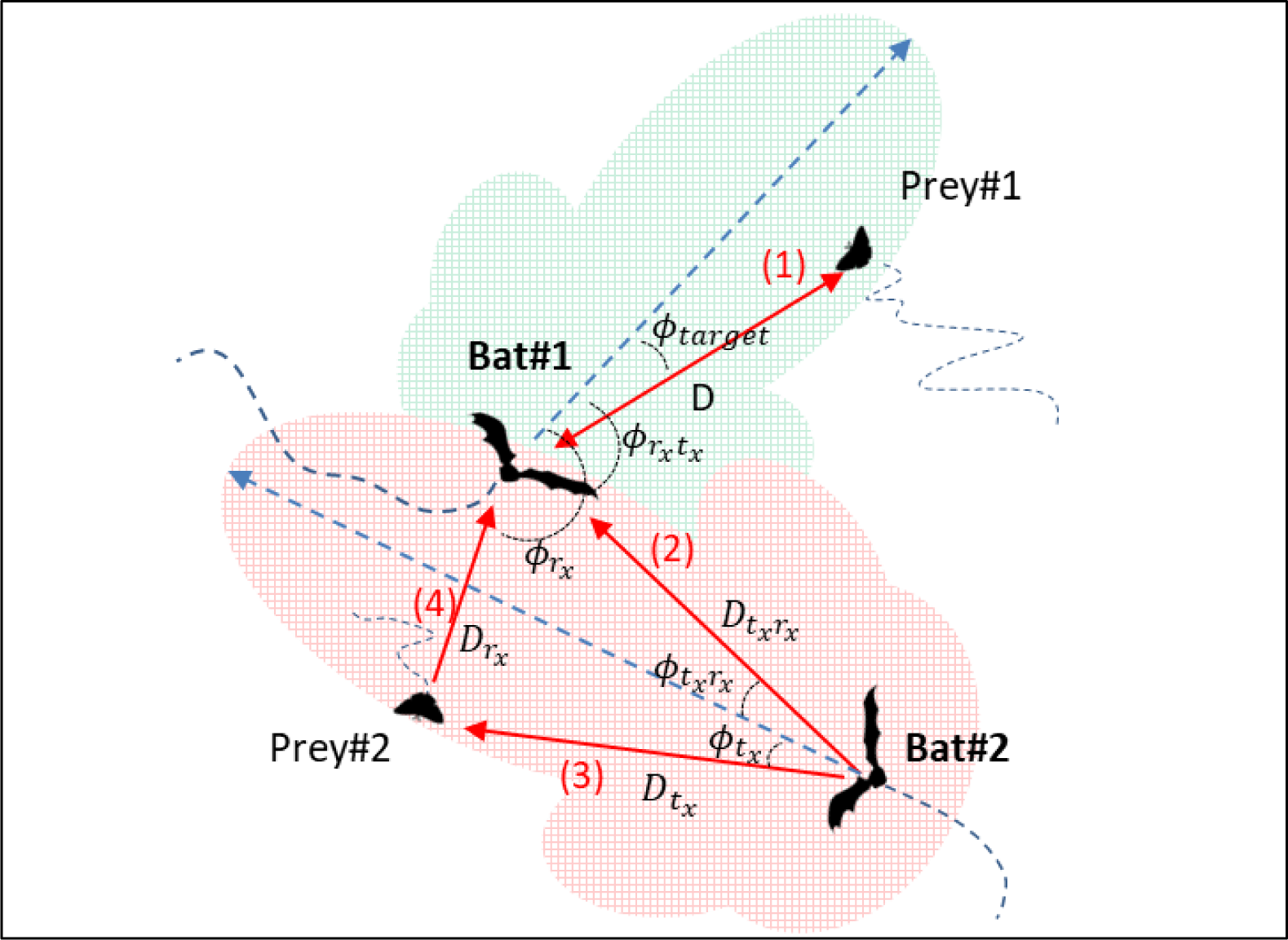
Angles and distances for two bats and two prey items. Bat#1 receives a reflected echo from prey#1 located at distance D, and angle Ф_target_ relative to its flight direction (arrow 1). Prey#1 is within the detection range of Bat#1, depicted by the green shaded piston area. Bat#1 also receives masking sounds: the signals emitted by Bat#2, arrive at the ear of Bat#1 at an angle Ф_t_x_r_x__ between arrow 2 and Bat #1 flight-direction and from a distance of D_t_x_r_x__. The echolocation signals of Bat#2 are reflected by Prey#2, and received by Bat#1, as masking signals at an angle Ф_r_x__, between arrows 4 and Bat#1 flight direction.

**Fig S 12:**
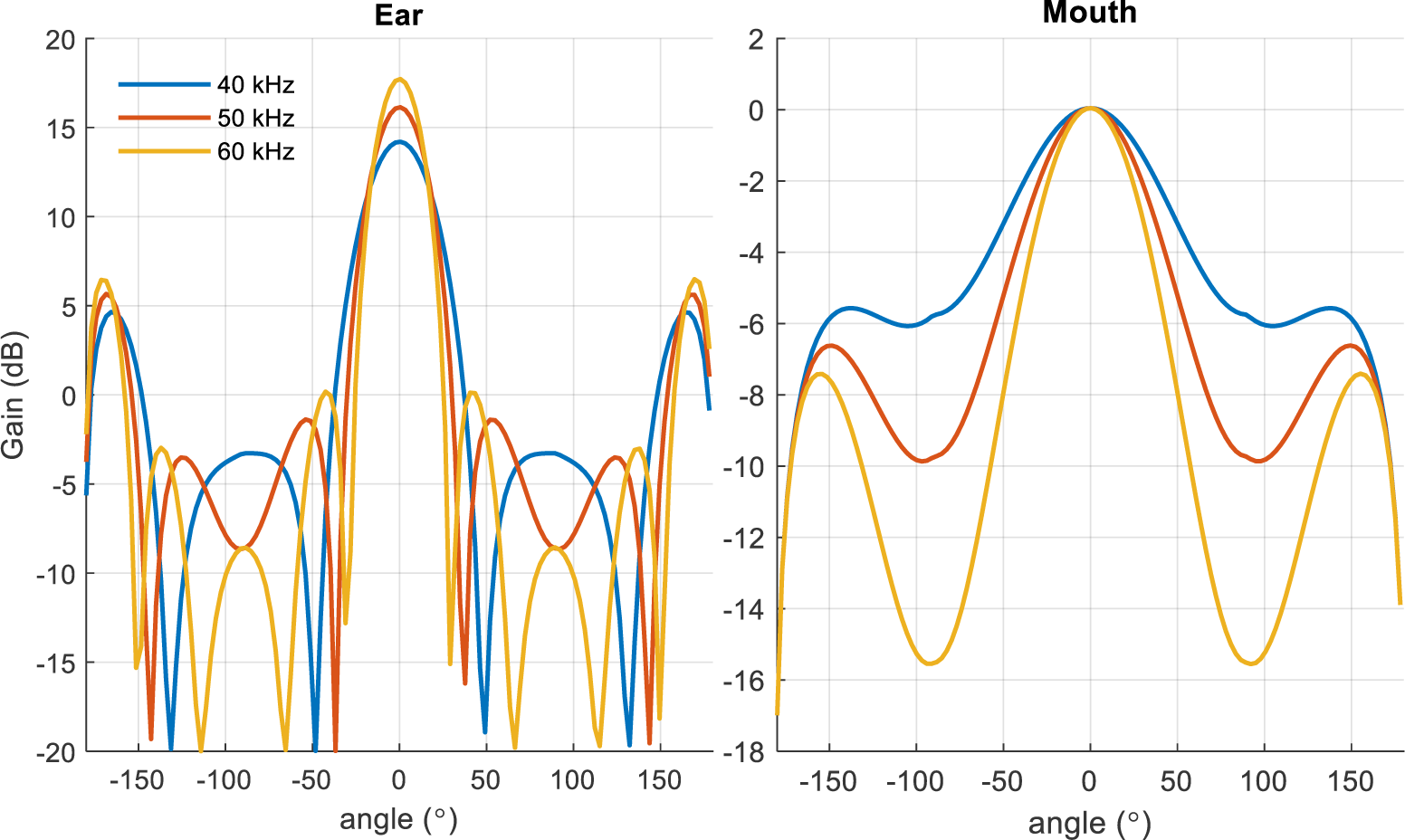
The modified piston model for the directivity of ear and mouth of the bat.

